# Lipocalin-2-expressing red pulp macrophages link ischemic stroke to systemic immune reprogramming in mice and humans

**DOI:** 10.64898/2026.06.23.733904

**Authors:** Alexandra Lucaciu, Patrick Wurzel, Stine Rytz Rasmussen, Eva Lueckhoff, Franziska Mayser, Jiffin Benjamin, Roxane-Isabelle Kestner, Verena Haas, Lisa Sophie Huber, Deepthi Bevara, Nico Landvogt, Melanie Glueck, Karen Gertz, Raphael Raspe, Sethuraman Subramanian, Christoph Welsch, Julia Bein, Peter J. Wild, Helena Radbruch, Christian Grefkes, Adam Strzelczyk, Waltraud Pfeilschifter, Michael H. Sieweke, Josef M. Pfeilschifter, Julien Subburayalu, Rajkumar Vutukuri

**Author notes:** These authors contributed equally and share last authorship (Julien Subburayalu, Rajkumar Vutukuri). Correspondence: Rajkumar Vutukuri, Ph.D.

## Abstract

**Background:** Acute ischemic stroke (AIS) triggers profound systemic immune alterations that contribute to post-stroke infections. The spleen is a key peripheral immune organ affected by AIS, but the molecular mediators linking cerebral ischemia to splenic immune dysfunction remain incompletely understood. Lipocalin-2 (LCN2) is an acute-phase protein implicated in neuroinflammation and host defense, yet its peripheral regulation, cellular sources and immune-modulatory actions after AIS remain unclear.

**Methods:** Male, 10-12-week-old C57BL/6J mice underwent transient middle cerebral artery occlusion (tMCAO) or sham surgery. Spleens were collected 24 hours, 3 days, or 7 days post-stroke for 3’-MACE-Seq transcriptomics, multiplex cytokine profiling, RT-qPCR, immunoblotting, and immunofluorescence. Murine T cells and monocytes were magnetically isolated, stimulated with recombinant LCN2 protein or vehicle, and analyzed using (i) RT-qPCR of LPS-induced proinflammatory cytokines (*Il1b*, *Il6*, *Tnf*), (ii) LCN2-driven T cell chemotaxis, (iii) FACS-based phenotyping, (iv) FACS-based phagocytosis of pHrodo-labeled *E*. *coli* bioparticles, and (v) FACS-based quantification of reactive oxygen species (ROS) using DHR-123. Human plasma samples and post-mortem spleens from three patients with ischemic stroke and one non-stroke control were analyzed for LCN2 expression and cellular localization. Untouched human monocytes from healthy donors were exposed to recombinant LCN2 and assessed by flow cytometry for antigen-presentation, co-stimulatory markers, phagocytosis and ROS production.

**Results:** Transcriptomics revealed robust, biphasic splenic *Lcn2* upregulation at 24 hours and 7 days after tMCAO. LCN2 localized predominantly to F4/80^+^ red pulp macrophages (RPMs), and accumulated at RPM-associated CD3^+^ contact sites. AIS induced a broad suppression of splenic innate and adaptive cytokine networks. Recombinant LCN2 induced T cell hyporesponsiveness, suppressed *Il6* and *Tnf* induction, and impaired CCL5-directed chemotaxis. In monocytes, LCN2 promoted a shift toward Ly6C^low^ phenotypes, enhanced phagocytosis of *E*. *coli*, and uncoupled phagocytosis from ROS production. In human ischemic stroke spleens, LCN2 was detected in CD68^+^ RPMs and at RPM-associated immune-cell contact sites, consistent with the murine splenic phenotype. In line with the murine findings, human monocytes exposed to LCN2 displayed increased *E*. *coli* uptake but reduced ROS, alongside reduced HLA-DR, CD80, CD206, and CD11b expression.

**Conclusions:** AIS induces a conserved splenic LCN2-expressing RPM niche in mice and humans. Recombinant LCN2 is sufficient to reprogram T cells and monocytes toward hyporesponsive, tolerogenic and phagocytosis-ROS-uncoupled states, thereby linking cerebral ischemia to systemic immune dysfunction. These findings identify LCN2-expressing RPMs as a translationally relevant immunoregulatory niche after AIS and nominate receptor-specific LCN2 signaling as a candidate pathway for future studies aimed at mitigating post-stroke immunodepression and infection susceptibility without compromising antimicrobial host defense.

## Introduction

Acute ischemic stroke (AIS) is a leading cause of death and disability worldwide, affecting over 13 million people annually, with few therapeutic options beyond recanalization (<u>Campbell et al. 2019</u>; <u>Sharma et Lee 2025</u>) (1,2). Although mechanical thrombectomy and thrombolysis have strongly improved acute care, neurological improvements often remain limited, especially for patients with large vessel occlusions or outside therapeutic windows (<u>Saver et al. 2016</u>) (3). Consequently, research increasingly targets secondary injury mechanisms to preserve the penumbra and to improve recovery (<u>Tymianski et al. 2025</u>; <u>Tymianski 2024</u>; <u>Hill et al. 2025</u>) (4–6).

The post-stroke immune response is pivotal for clinical outcomes, marked by an initial neuroinflammatory phase followed by profound systemic immunosuppression (<u>Iadecola et Anrather 2011</u>; <u>Meisel et al. 2005</u>) (7,8). This central nervous system (CNS) injury-induced immunodepression (CIDS) manifests as lymphocytopenia, impaired antigen presentation, and deficits in innate and adaptive immunity, predisposing patients to infections (<u>Prass et al. 2003</u>; <u>Westendorp et al. 2011</u>; <u>Dirnagl et al. 2007</u>) (9–11). The extent of CIDS correlates with stroke severity and predicts poor outcomes and mortality (<u>Vogelgesang et al. 2008</u>; <u>Klehmet et al. 2009</u>; <u>Vaghi et al. 2024</u>) (12–14). Despite its clinical significance, effective strategies to counteract CIDS are lacking, highlighting the need for molecular elucidation.

The spleen is a central orchestrator of systemic immune responses following stroke (<u>Ajmo et al. 2008</u>; <u>Seifert et al. 2012</u>; <u>Dotson et al. 2016</u>) (15–17). After AIS, the spleen undergoes structural and functional changes, including atrophy, immune cell depletion, and altered macrophage phenotypes (<u>Offner et al. 2006a</u>; <u>Offner et al. 2006b</u>; <u>Vendrame et al. 2006</u>) (18–20). These changes are driven by an activation of the hypothalamic-pituitary-adrenal axis and the sympathetic nervous system, which reprogram splenic immune cells before their recruitment to the injured CNS (<u>Mracsko et al. 2014</u>; <u>Mcculloch et al. 2017</u>) (21,22). Thus, the spleen acts as both a reservoir and conditioning center for immune cells destined for cerebral recruitment, directly influencing neuroinflammation and stroke outcomes (<u>Ajmo et al. 2008</u>; <u>Swirski et al. 2009</u>) (15,23).

Lipocalin-2 (LCN2), a 25-kDa glycoprotein, acts as a key mediator of neuroinflammation and immune reprogramming in stroke (<u>Jha et al. 2015</u>; <u>Jin et al. 2014</u>) (24,25). LCN2 is rapidly upregulated in brain and peripheral tissues after CNS injury, functioning as an acute-phase and iron-sequestering protein (<u>Berard et al. 2012</u>; <u>Dekens et al. 2018</u>) (26,27). In the CNS, LCN2 expression by reactive astrocytes and microglia promotes neuronal death via iron-mediated oxidative stress (<u>Bi et al. 2013</u>; <u>Guo et al. 2025</u>) (28,29), and disrupts the blood-brain barrier (BBB) by stabilizing MMP-9 (<u>Yan et al. 2001</u>) (30). Elevated plasma LCN2 levels correlate with stroke severity, infarct volume, and poor outcomes in preclinical and clinical studies (<u>Huang et al. 2025</u>; <u>Hochmeister et al. 2016</u>) (31,32).

Beyond the CNS, LCN2 is a critical immune modulator in peripheral lymphoid organs. Internalization via 24p3R (SLC22A17) and megalin (LRP2) triggers signaling cascades affecting immune cell activation, migration, and function (<u>Devireddy et al. 2005</u>; <u>Hvidberg et al. 2005</u>; <u>Yang et al. 2002</u>) (33–35). Short-term LCN2 engagement promotes proinflammatory cytokine production in macrophages (<u>Cheng et al. 2015</u>; <u>Oberoi et al. 2015</u>) (36,37), while sustained exposure has been linked to impaired T cell activation, T cell hyporesponsiveness, and monocyte tolerance via IGF-1R signaling (<u>Warszawska et al. 2013</u>; <u>Watzenboeck et al. 2021</u>; <u>Czech et al. 2024</u>) (38–40), potentially explaining CIDS.

Despite increasing recognition of LCN2 as a circulating biomarker after stroke, it remains unclear whether splenic LCN2 expression reflects passive systemic inflammation or a spatially organized immunoregulatory niche within peripheral lymphoid tissue. This distinction is clinically important because circulating LCN2 measurements cannot resolve cellular source, tissue compartment or immune-cell interactions. Moreover, LCN2 has context-dependent roles in host defense: it can restrict selected bacterial infections through siderophore sequestration (<u>Nairz et al. 2015</u>; <u>Chan et al. 2009</u>; <u>Moschen et al. 2016</u>; <u>Zhang et al. 2023</u>; <u>Wu et al. 2010</u>; <u>Cramer et al.</u> <u>2017</u>) (41–46), but can also deactivate macrophages and modulate antibacterial or antiviral immunity in a pathogen– and compartment-specific manner (<u>Warszawska et al. 2013</u>; <u>Watzenboeck et al. 2021</u>) (38,39). Thus, defining the splenic cellular source and functional consequences of LCN2 after AIS is required before LCN2 signaling can be considered as a therapeutic axis.

Here, we show that AIS triggers biphasic splenic LCN2 upregulation that localizes predominantly to red pulp macrophages (RPMs). By integrating temporal splenic transcriptomics, spatial immunofluorescence, murine immune-cell perturbation assays, human plasma analyses, postmortem human ischemic stroke spleen tissue and functional human monocyte assays, we identify a conserved splenic LCN2-expressing RPM axis that links cerebral ischemia to peripheral immune reprogramming. Our data support a model in which LCN2 is sufficient to induce T cell and monocyte hyporesponsiveness, including uncoupling of *E*. *coli* particle uptake from reactive oxygen species (ROS) production, thereby providing a mechanistic framework for future studies of post-stroke infection susceptibility.

## Results

### Transcriptomic profiling reveals robust splenic *Lcn2* upregulation and immune pathway activation after acute ischemic stroke

To elucidate the transcriptomic signature of the spleen in response to acute cerebral ischemia, we performed 3’-MACE-Seq of RNA isolated from spleens of C57BL/6J mice subjected to transient middle cerebral artery occlusion (tMCAO) for 1 hour or sham surgery with subsequent analysis after 24 hours (n = 3 per group; Fig. 1A). Principal component analysis (PCA) of normalized gene expression profiles revealed distinct separation between tMCAO and sham samples, with PC1 and PC2 accounting for 47.81% and 14.93% of the variance, respectively, underscoring the robust impact of ischemic injury on the splenic transcriptome (Fig. 1B). Differential expression analyses identified 347 up-regulated and 564 down-regulated genes (adjusted *P* < 0.001, log2FC > |2.0|; Fig. 1C), reflecting a broad and coordinated transcriptional response to tMCAO.

**Figure 1.**
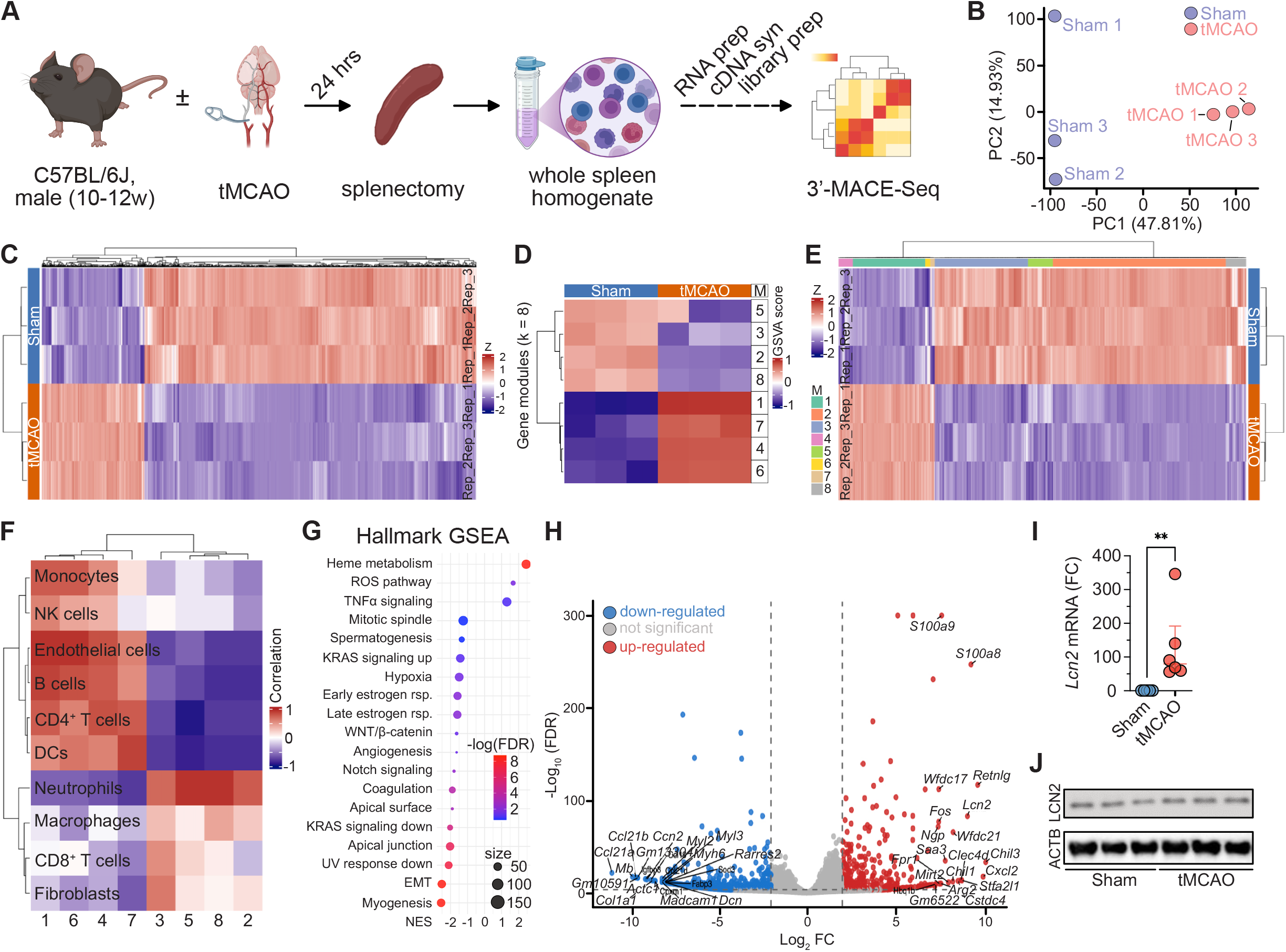
*Lcn2* is up-regulated in the spleen in the acute phase of ischemic stroke. (**A**) Experimental workflow: Male, 10-12 week-old C57BL/6J mice underwent transient middle cerebral artery occlusion (tMCAO) or sham surgery. After 24 hours, spleens were harvested, homogenized, and analyzed by 3’-MACE-Seq. (**B**) Principal component analysis (PCA) shows clear separation of global gene expression profiles between tMCAO-(orange) or sham (blue) spleens (three independent experiments). (**C**) Heatmap of the top 1,000 differentially expressed genes (adjusted *P* ≤ 0.001). (**D**) Unsupervised k-means clustering identifies eight gene modules (M1-M8), visualized by GSVA scores (−1 to +1). (**E**) Same data as in (**C**), reorganized according to modules in (**D**). (**F**) Pearson correlation heatmap of gene modules with immune cell signatures using manual ssGSEA based on canonical markers for neutrophils, macrophages, monocytes, NK cells, CD8^+^ T cells, CD4^+^ T cells, B cells, dendritic cells, endothelial cells, and fibroblasts (marker selection shown in Table 1). (**G**) Gene set enrichment analysis (GSEA) of Hallmark pathways displays normalized enrichment scores (NES), gene counts, and false discovery rates (FDR) for significantly up– or down-regulated pathways. (**H**) Volcano plot showing 347 up-regulated (red) and 564 down-regulated genes (blue) in tMCAO versus sham spleens (adjusted *P* ≤ 0.001, log2FC ≥ | 2.0 |); the top 20 genes are labeled. (**I**) RT-qPCR validation of *Lcn2* mRNA expression normalized to *Gapdh* and expressed as fold-change (ΔΔCT) in sham (blue) and tMCAO (red) spleens (*n* = 5-6 per group, pooled from six experiments; median ± IQR; Mann-Whitney test; \*\**P* ≤ 0.01). (**J**) Immunoblot of LCN2 protein relative to β-Actin (ACTB) in spleen lysates from sham and tMCAO mice (three independent experiments).

**Table 1.**
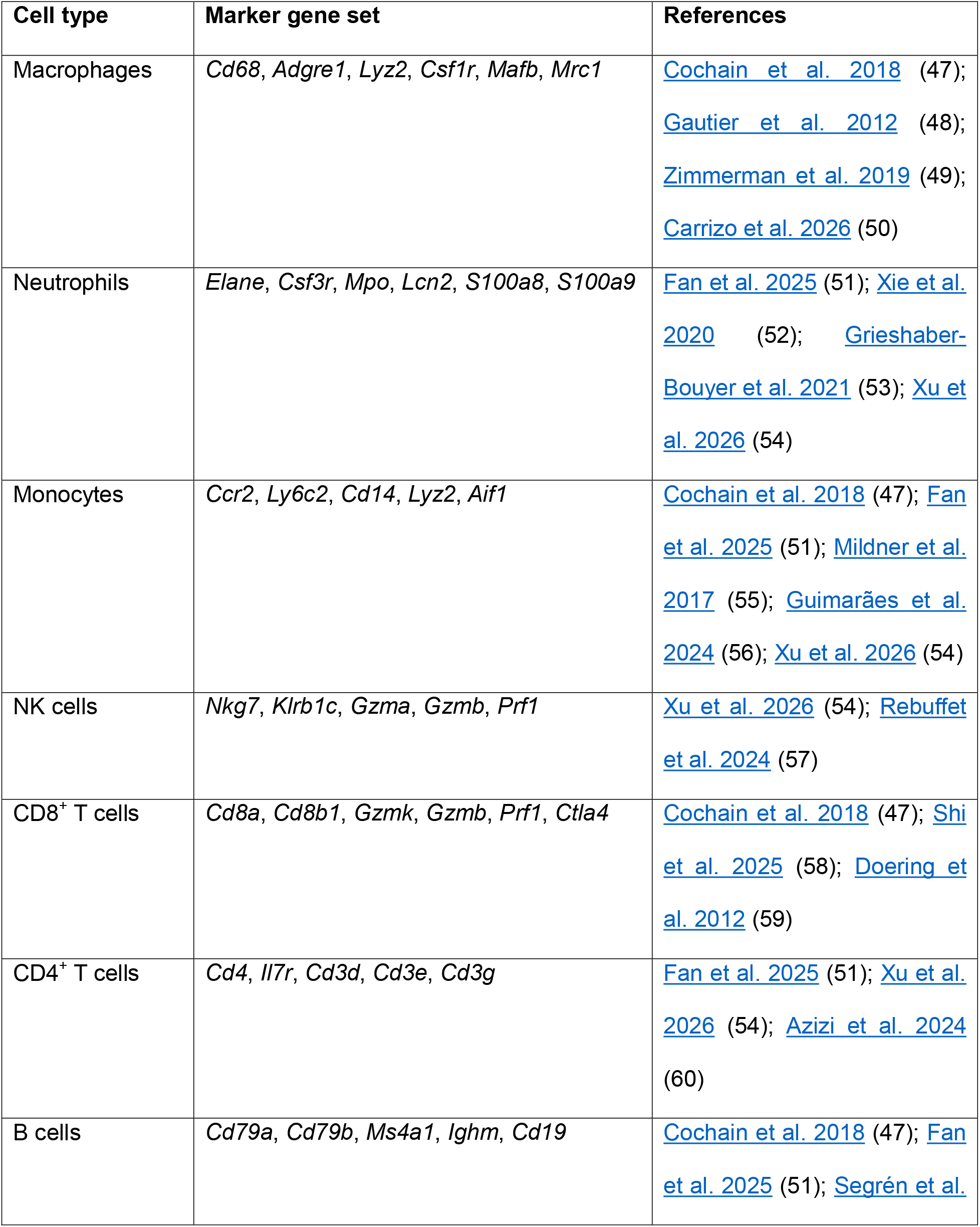
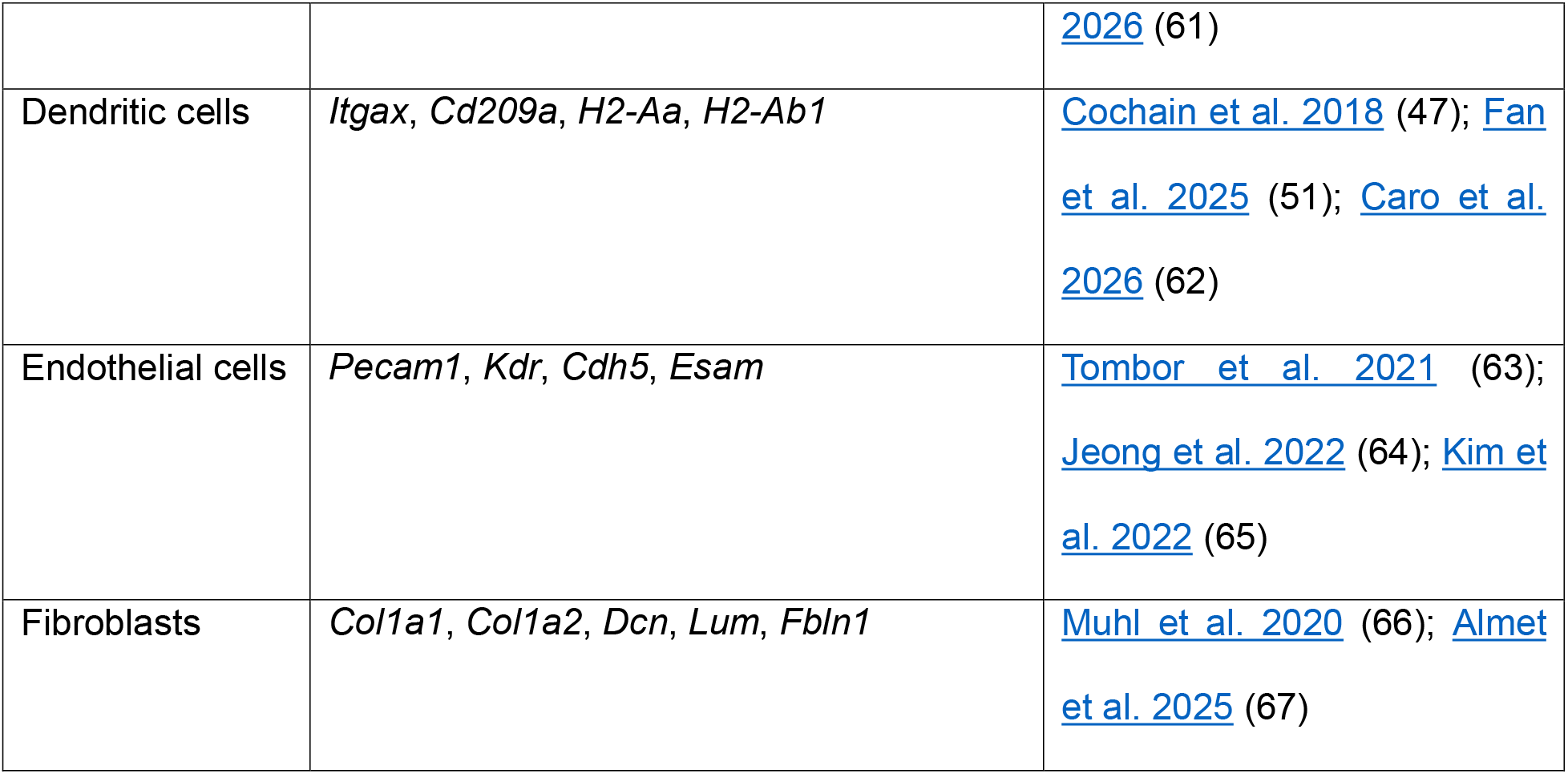
Curated murine cell type-associated marker gene sets used for bulk 3’-MACE-Seq-derived transcriptomic module correlation analysis.

To further dissect the changes of the splenic response, we performed an unsupervised hierarchical clustering of differentially expressed genes (DEGs), thereby identifying eight distinct gene modules (k = 8), with module-specific gene set variation analysis (GSVA) scores highlighting divergent module activation between sham and tMCAO samples (Fig. 1D). This analysis allowed us to cluster the top 1000 DEGs (adjusted *P* < 0.001) according to the identified gene modules (Fig. 1E), and to detect marker genes of each cluster (Supplementary Figure 1A-H). Subsequently, we performed a correlation analysis of those genes contained within each module with established canonical and (immune) cell type-associated marker gene sets (Table 1) to identify associations between the identified gene modules with (immune) cell subtypes of the spleen, including macrophages (47–50), neutrophils (51–54), monocytes (47,51,54–56), NK cells (54,57), CD8^+^ T cells (47,58,59), CD4^+^ T cells (51,54,60), B cells (47,51,61), dendritic cells (47,51,62), endothelial cells (63–65), or fibroblasts (66,67). This analysis revealed that the observed transcriptomic reprogramming was closely linked to shifts in transcriptional immune cell activation in the spleen (Fig. 1F).

To identify the biological pathways underlying these changes, we performed a comprehensive gene set enrichment analysis (GSEA) using several curated databases (Fig. 1G; Supplementary Fig. 2A-D). The GSEA was performed on pre-ranked gene lists using the Broad Institute’s GSEA tool; significantly enriched pathways were defined by false-discovery rate (FDR; *Q* < 0.1 and NES > |1.0|). The strongest up-regulated pathways included “Heme metabolism”, “ROS pathway”, and “TNFα signaling” (Fig. 1G, Table 2). Conversely, down-regulated pathways included “EMT”, “KRAS signaling down”, “WNT/β-catenin”, and “apical junction”, suggesting a pronounced activation of inflammatory and metabolic programs alongside suppression of endothelial or structural integrity of the spleen following ischemic injury (Fig. 1G). These enrichment patterns were consistent across multiple databases, with immune activation pathways prominently expressed (Supplementary Fig. 2A-D).

**Table 2.** GSEA of significantly altered pathways 24-hours tMCAO vs. sham using the Hallmark database.

| ID_Description | NES | Q value |
| --- | --- | --- |
| HALLMARK_HEME_METABOLISM | 2.43373609 | 1.6667E-09 |
| HALLMARK_REACTIVE_OXYGEN_SPECIES_PATHWAY | 1.70580267 | 0.03066158 |
| HALLMARK_TNFA_SIGNALING_VIA_NFKB | 1.303946 | 0.09334757 |
| HALLMARK_KRAS_SIGNALING_UP | -1.3962384 | 0.08013101 |
| HALLMARK_HYPOXIA | -1.469382 | 0.02732193 |
| HALLMARK_ESTROGEN_RESPONSE_EARLY | -1.5725899 | 0.01057278 |
| HALLMARK_ESTROGEN_RESPONSE_LATE | -1.5832914 | 0.00629688 |
| HALLMARK_ANGIOGENESIS | -1.6178944 | 0.04480036 |
| HALLMARK_WNT_BETA_CATENIN_SIGNALING | -1.6200684 | 0.04682391 |
| HALLMARK_NOTCH_SIGNALING | -1.7737733 | 0.00634109 |
| HALLMARK_COAGULATION | -1.8563762 | 0.00010534 |
| HALLMARK_APICAL_SURFACE | -1.873248 | 0.00120559 |
| HALLMARK_APICAL_JUNCTION | -1.994432 | 3.1803E-07 |
| HALLMARK_KRAS_SIGNALING_DN | -2.0053071 | 9.0332E-06 |
| HALLMARK_UV_RESPONSE_DN | -2.0654432 | 2.0425E-08 |
| HALLMARK_EPITHELIAL_MESENCHYMAL_TRANSITION | -2.4752053 | 1.6667E-09 |
| HALLMARK_MYOGENESIS | -2.5161573 | 1.6667E-09 |

Interrogation of the most abundantly regulated genes revealed that *Lcn2* was among the top upregulated transcripts in tMCAO spleens (Fig. 1H, Supplementary Table 1). To validate these findings and gauge functional relevance of *Lcn2* as a potential immunomodulatory mediator (<u>Cheng et al. 2015</u>; <u>Oberoi et al. 2015</u>) (36,37), we performed a real-time, quantitative polymerase chain reaction (RT-qPCR) and an immunoblot analysis on splenic lysates. Indeed, RT-qPCR confirmed robust transcriptional up-regulation of *Lcn2* following tMCAO (*P* < 0.001; Fig. 1I), whilst the immunoblot analysis corroborated increased LCN2 protein expression in tMCAO spleens (Fig. 1J).

In summary, these data demonstrate that AIS triggers a robust transcriptomic reprogramming of the spleen, characterized by activation of inflammatory and metabolic pathways, and suppression of gene programs related to splenic structural integrity, with *Lcn2* emerging as one of the most prominently regulated genes.

### Temporal splenic transcriptomics reveal biphasic immune remodeling after acute ischemic stroke

Building on the acute 24-hour splenic transcriptomic profiling that identified *Lcn2* as one of the most strongly induced genes after tMCAO, we next sought to define the temporal architecture of splenic immune remodeling after AIS. To this end, we performed 3’-MACE-Seq on spleens harvested 3 days and 7 days after tMCAO and undertook a comparative analysis (Supplementary Table 2). PCA revealed that 24-hour and 7-day post-tMCAO spleens formed distinct clusters separated from sham controls, whereas 3-day post-tMCAO samples were positioned closer to sham on PC1 but remained separated on PC2, suggesting a transient phase of relative transcriptomic quiescence rather than complete normalization (Fig. 2A). Consistent with this interpretation, differential expression analysis identified only limited transcriptional divergence at day 3, while day 7 spleens again displayed broad immune and inflammatory gene regulation resembling the acute post-stroke phase (Fig. 2B,C; Supplementary Fig. 3A-F). Among the most strongly up-regulated transcripts at day 7 were innate immune, neutrophil-associated and antimicrobial genes, including *S100a8*, *Lcn2*, *Elane*, *Retnlg*, *Chil3*, *Hbb-bs*, and *Hba-ps3* (<u>Ryckman et al. 2003; Huang et al. 2017</u>) (68,69), indicating delayed re-engagement of host-defense and acute-phase programs after AIS (Fig. 2C).

**Figure 2.**
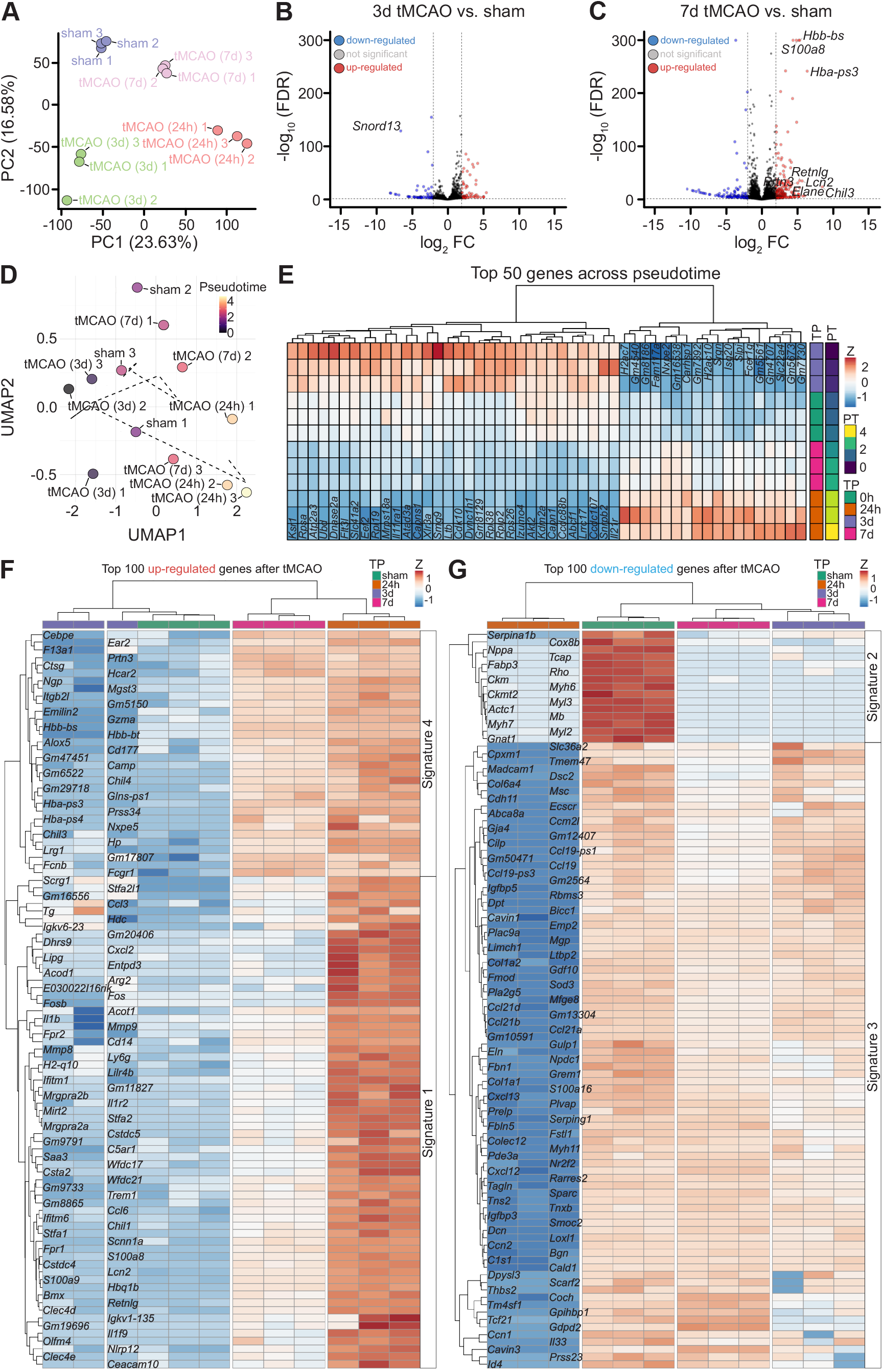
Dynamic transcriptomic reprogramming of the spleen from the acute to chronic phases following ischemic stroke. (**A**) Principal component analysis (PCA) of global gene expression reveals distinct clustering of splenocytes isolated from different time-points after tMCAO-(24 hours: red; 3 days: green; 7 days: pink) or sham-operated (blue) C57BL/6J mice, from three independent experiments. (**B**) Volcano plot of the differentially expressed genes (DEGs) between sham-operated or 3 days after tMCAO in splenocytes isolated from C57BL/6J mice, showing up-regulated (red) and down-regulated genes (blue, *P*-adjusted ≤ 0.001, log2FC ≥ | 2.0 |). (**C**) Same as (**B**) but comparison against splenocytes isolated from C57BL/6J mice 7 days after tMCAO. (**D**) Uniform manifold approximation and projection (UMAP) plot showing global gene expression profiles across pseudotime of 3’-MACE-Seq of splenocytes isolated from tMCAO-(following different timepoints) or sham-operated (blue) C57BL/6J mice, from three independent experiments. (**E**) Top 50 genes from (**D**) showing positive or negative regulation across pseudotime (PT), illustrating the temporal evolution of gene expression from the timepoints (TP) following tMCAO including the acute (0 to 24 hours) to subacute (3 days) and chronic phases (7 days) after tMCAO as compared to sham-operated splenocytes isolated from C57BL/6J mice, from three independent experiments. Each row represents a gene, and each column represents a sample ordered by PT. (**F**) Gene expression profile of the top 100 up-regulated genes, thereby identifying signature genes whose expression is recovered to baseline after an initial strong expression 24 hours after tMCAO (“Signature 1”) and such that are maintained up at 24 hours and 7 days following tMCAO (“Signature 4”) as compared to sham-treated splenocytes. (**G**) Same as (**F**) but showing the top 100 down-regulated genes in splenocytes isolated from tMCAO– or sham-operated (green) C57BL/6J mice kinetically tracked across the different timepoints (TP) after tMCAO (3 days: violet; 7 days: pink), from three independent experiments. The heatmap uncovers several genes whose expression is recovered at day 3 and day 7 following tMCAO (“Signature 3”) and such that are persistently down-regulated (“Signature 2”) as compared to sham-treated splenocytes.

To further resolve this temporal pattern, we applied a UMAP-based pseudotime analysis across all splenic transcriptomes. Sham and 3 days post-tMCAO samples occupied earlier pseudotime states, whereas 24-hour and 7 days post-tMCAO samples shifted toward later pseudotime states, supporting a coordinated transition from baseline or relatively quiescent splenic states toward acute and delayed inflammatory states (Fig. 2D). Genes increasing along pseudotime were enriched for immune activation and effector functions (e.g., *Srgn*, *Isg20*, *Slpi*, *Fcer1g*, *Slc22a4*) (<u>Weiss et al. 2018</u>; <u>Huang et al. 2024</u>; <u>Tyshchenko et al. 2025</u>) (70–72), whereas genes decreasing along pseudotime included transcripts associated with protein synthesis, metabolic regulation (e.g., *Ksr1*, *Rpsa*, *Atp2a3*, *Eef2*, *Rpl19*, *Mrps18a*, *Il11ra1*) (<u>Bell et al. 1999</u>; <u>Nagle et al. 2024</u>; <u>Kaul et al. 2011; Bee et al. 2011</u>) (73–76), and structural programs (Fig. 2E). Thus, global trajectory analysis supports a biphasic model of splenic immune remodeling after AIS, characterized by early inflammatory activation, partial subacute attenuation, and delayed reactivation of immune and host-defense pathways.

To refine this temporal architecture, we next tracked the top acutely regulated genes across all time points and identified four major temporal gene-expression signatures (Fig. 2F,G). First, a transient acute-response signature (signature 1) comprised genes that peaked at 24 hours and were attenuated thereafter, including inflammatory and chemotactic mediators such as *Il1b*, *Fos*, *Ccl3*, *Retnlg*, *Cd14*, and *C5ar1*. This signature was enriched for immune response-regulating receptor signaling, response to LPS, chemotaxis, interleukin-1 response and neutrophil chemotaxis (Supplementary Fig. 3G), consistent with rapid innate immune activation after stroke. Second, a maintained-down signature (signature 2) contained structural, cytoskeletal and metabolic genes, including genes associated with muscle contraction, actin organization, and arginine/proline metabolism (Supplementary Fig. 3H), indicating persistent suppression of non-immune structural and metabolic programs in the spleen, aligning with the concept of resource allocation away from non-immune functions during systemic inflammatory stress (<u>Wang et al. 2025</u>) (77). Third, a recovered-up signature (signature 3) comprised genes that were acutely suppressed but returned toward baseline at later time points, including genes related to extracellular matrix organization, cell adhesion and tissue remodeling (Supplementary Fig. 3I). Fourth, a maintained or re-emergent up-regulated signature (signature 4) including host-defense and antimicrobial genes such as *Fcgr1*, *Lrg1*, *Chil3*, *Hbb-bs*, *Camp*, and *Elane*, with enrichment for neutrophil-associated, antimicrobial and infection-related pathways (Supplementary Fig. 3J), consistent with prior reports of early immune suppression followed by recovery in peripheral organs after stroke (<u>Iadecola et al. 2020</u>) (78).

Within this temporal framework, *Lcn2* displayed a biologically notable pattern. It was among the most strongly induced acute-phase genes at 24 hours, was attenuated during the 3 day relative quiescence phase, and reappeared among highly regulated immune effector transcripts at day 7. Together, these temporal transcriptomic analyses identify AIS-induced splenic remodeling as a dynamic, phase-specific process and nominate *Lcn2* as a priority candidate linking cerebral ischemia to delayed peripheral immune reprogramming.

### LCN2-expressing red pulp macrophages define a splenic immunomodulatory niche after acute ischemic stroke

Following our identification of temporally distinct splenic gene expression signatures after tMCAO, we next sought to define the immunological context of *Lcn2*, one of the most robustly up-regulated genes in the acute and delayed post-stroke phases. Given its dynamic regulation and association with immune and host-defense transcriptional programs, we asked whether the *Lcn2*^high^ splenic state was accompanied by altered cytokine networks and whether LCN2 localized to a defined cellular niche within the spleen. To address this, we first performed a comprehensive 40-plex cytokine profiling using spleen protein-homogenates from sham and tMCAO-treated mice at 24 hours, 3 days, and 7 days post-injury (Fig. 3A). This analysis revealed a broad reduction of cytokines pertaining to both the adaptive and innate immune cell lineage as compared to sham-treated controls (Fig. 3B; Supplementary Fig. 4A,B). The global down-regulation of cytokines relevant for innate and adaptive immune responses is consistent with the well-documented phenomenon of post-stroke immunodepression, which has been linked to increased infection risk and impaired immunosurveillance in both clinical and experimental settings (<u>Westendorp et al. 2011</u>; <u>Liesz et al. 2009</u>) (10,79). Given the prominent up-regulation of *Lcn2* in our transcriptomic data, we next validated its expression and characterized its spatial distribution using RT-qPCR, immunoblotting, and immunofluorescence (IF) across all time points post-injury (Fig. 3C). Here, *Lcn2* mRNA and protein levels were transiently reduced at day 3 post-tMCAO but showed a marked increase by day 7 similar to 24 hours post-injury (Fig. 3D,E). This biphasic regulation suggests that *Lcn2* transcription is tightly controlled in the spleen, and may thus contribute to both immunosuppressive and immunostimulatory pathways. These findings are in line with recent reports demonstrating that LCN2 acts as a dynamic acute-phase protein in both central and peripheral immune compartments following CNS injury. Thus, LCN2 may serve as a molecular switch between phases of immunosuppression and immune reactivation (<u>Guo et al. 2025</u>; <u>Xie et al. 2023</u>; <u>Mao et al. 2016</u>; <u>Saenz-Pipaon et al. 2023</u>; <u>Wang et al. 2024</u>) (29,80–83). To further dissect the spatial and cellular context of LCN2 expression, we performed immunofluorescence staining on spleen sections (Fig. 3F). Qualitative and quantitative image analysis revealed a significant increase of LCN2 expression selectively in the red pulp, which we could allocate to myeloid cells, especially F4/80^+^ red-pulp macrophages (RPMs) (Fig. 3F,G), which increased over time (Fig. 3G). Conversely, the number of red pulp CD3^+^ T cells declined markedly at days 3 and 7 (Fig. 3G), aligning with previous reports on splenic T cell egress following AIS (<u>Lucaciu et al. 2020</u>) (84) and recent findings of an inverse correlation of CD3^+^ T cells within LCN2-enriched micromilieus such as tumor microenvironments (TME) (<u>Bossowski et al. 2026</u>) (85). In fact, image analyses revealed that LCN2-expressing RPMs colocalized at sites of close contact with CD3^+^ T cells, consistent with the formation of cell-to-cell contact sites (Fig. 3H). These qualitative findings support a model in which RPMs represent a major splenic source and spatial hub of LCN2-associated immune modulation after stroke. The observed reduction in red pulp CD3^+^ T cell area, coinciding with increased LCN2 expression and macrophage-T cell contact sites, suggests that LCN2-expressing RPMs may influence T cell retention, egress, or activation status in the splenic microenvironment. This is consistent with emerging literature demonstrating that LCN2 can modulate both innate and adaptive immune responses, including polarization of circulating monocytes or macrophages, regulation of T cell function, and shaping the local cytokine milieu (<u>Sciarretta et al. 2023</u>; <u>Xu et al. 2023</u>) (86,87). Mechanistically, LCN2 has been shown to act through its receptor 24p3R, which is expressed on multiple immune cell types, including neutrophils, monocytes, macrophages, dendritic cells, and T cells (<u>Berard et al. 2012</u>; <u>Czech et al. 2024</u>; <u>Chen et al. 2020</u>; <u>Jha et al. 2014</u>) (26,40,88,89). In the context of cerebral ischemia, LCN2 may promote neuroinflammation by amplifying cytokine release and facilitating leukocyte infiltration, while in peripheral tissues, it may serve as a critical link between central and systemic immune responses, orchestrating the balance between immunosuppression and re-activation. In summary, our data position LCN2-expressing RPMs at the core of splenic immunodynamics following cerebral ischemia.

**Figure 3.**
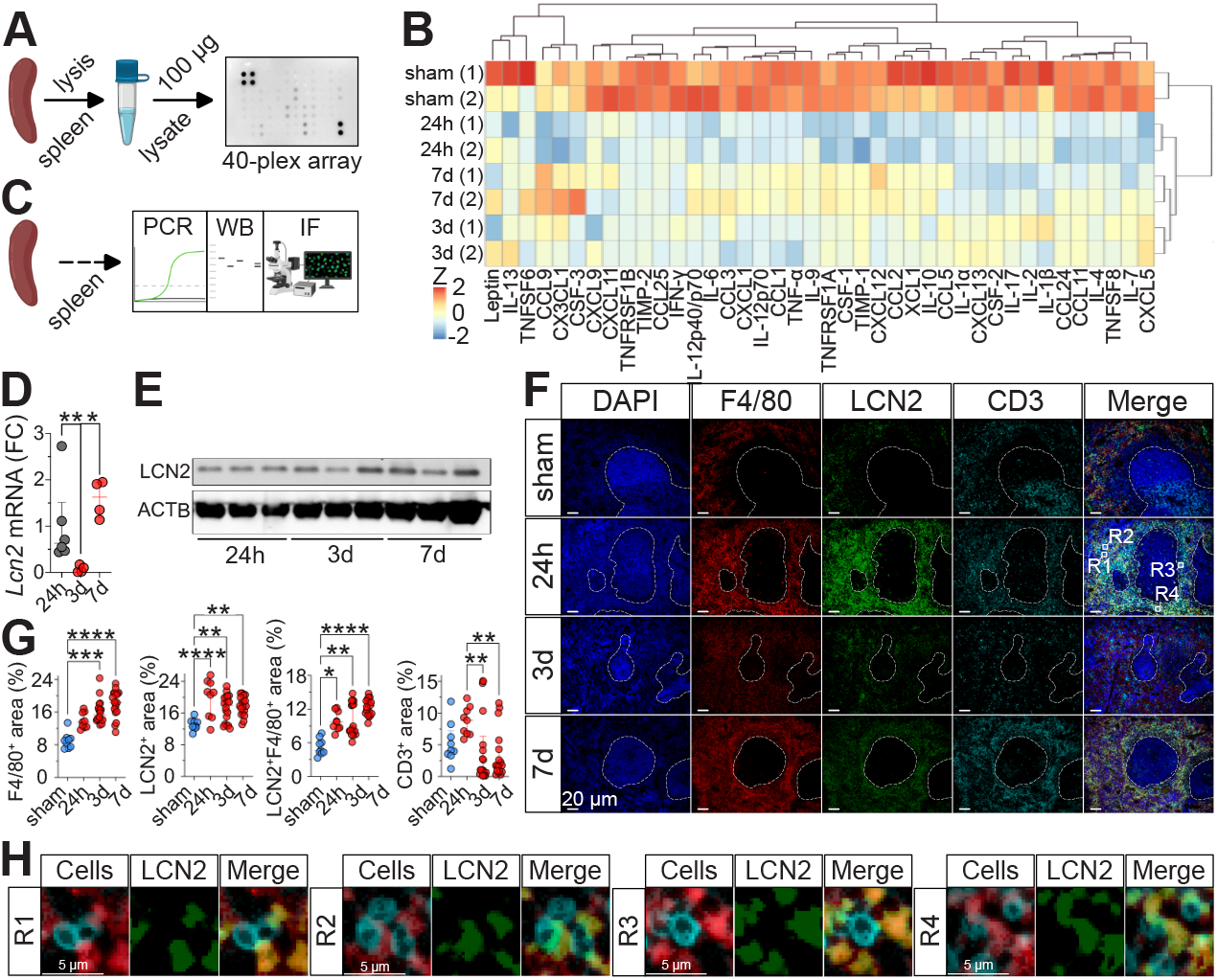
Lipocalin-2 is expressed in RPMs following ischemic stroke and contributes to cell-to-cell contact sites with T cells. (**A**) Experimental workflow: male, 10-12 week-old C57BL/6J mice underwent tMCAO or sham surgery. At 24 hours, 3 days, or 7 days, spleens were harvested and 100 µg lysate analyzed using a 40-plex inflammation antibody array. (**B**) Heatmap of cytokines and chemokines measured by multiplex immunoassay, clustered by treatment condition, analyzed in duplicate (each duplicate pooled from *n* = 4 mice). Representative of two independent experiments. (**C**) Similar design as in (**A**). Spleen lysates were used for RT-qPCR and immunoblotting, and FFPE sections for immunofluorescence (IF). (**D**) Relative *Lcn2* mRNA normalized to *Gapdh* and shown as fold-change (FC) at 24 hours (grey), 3 days, or 7 days (red). Data (*n* = 4-6 per group) are median ± IQR from up to six experiments; Mann-Whitney U test; \**P* ≤ 0.05, \*\**P* ≤ 0.01. (**E**) Immunoblot of LCN2 protein vs. β-Actin (ACTB) at indicated time points (three independent experiments). (**F**) Representative IF of spleen sections from sham or tMCAO mice across time points. Individual channels for DAPI, F4/80, LCN2, CD3, and the merged composite are shown. Scale bar = 20 µm. (**G**) Quantitative image analysis of red pulp regions from (**F**) showing area of F4/80, LCN2, LCN2-F4/80 double-positive, and CD3. Data are median ± IQR (*n* = 3-6 mice pooled per group; 3 ROIs per mouse). Kruskal-Wallis with Dunn’s test (F4/80, LCN2/F4/80, CD3) or one-way ANOVA with Tukey’s test (LCN2) were used. \**P* ≤ 0.05, \*\**P* ≤ 0.01, \*\*\**P* ≤ 0.001, \*\*\*\**P* ≤ 0.0001. (**H**) High-resolution IF images from regions R1-R4 in (**F**), showing F4/80^+^ RPMs (red) forming LCN2-rich (green) cell-to-cell contact sites with CD3^+^ T cells (turquoise). Scale bar = 5 µm.

### LCN2 is sufficient to induce hyporesponsive and phagocytosis-ROS-uncoupled phenotypes in splenic T cells and monocytes

Considering the discovery that LCN2-expressing RPMs orchestrate splenic immune modulation and T cell dynamics after AIS, we next investigated whether LCN2 exerts direct functional effects on isolated splenic T cells and monocytes previously shown to express the LCN2 receptor 24p3R (<u>Berard et al. 2012</u>; <u>Chen et al. 2020</u>) (26,88). This approach aimed to mechanistically link the *in vivo* immunomodulatory signatures to cell-intrinsic responses, thereby clarifying how LCN2 shapes the post-stroke immune landscape. To address this, we exposed splenic T cells and monocytes to recombinant LCN2 protein (Fig. 4A). Pre-incubation of T cells with LCN2 significantly reduced their capacity to up-regulate transcription of *Il6* and (of borderline significance) *Tnf* in response to LPS stimulation as compared to LPS alone (Fig. 4B). This hyporesponsive phenotype is consistent with the emerging paradigm that certain stimuli can induce a tolerogenic state in T cells, characterized by diminished proinflammatory cytokine production and impaired effector function, a phenomenon previously observed rather in models of sterile inflammation and infection (<u>Zajac et al. 1998</u>; <u>Mitchison 1964</u>; <u>Boomer et al. 2011</u>) (90–92). Mechanistically, binding of LCN2 via its receptor 24p3R can result in suppressed T cell activation and cytokine release via JAK-STAT and MAPK signaling (<u>Wang et al. 2021</u>; <u>Chen et al. 2026</u>) (93,94). To assess whether LCN2 also influenced T cell trafficking as suggested by diminished red pulp CD3^+^ T cells post-stroke (Fig. 3G), we performed a transwell migration assay using CCL5 (RANTES) as a migration-inducing chemoattractant (<u>Murooka et al. 2008</u>) (95). The addition of LCN2 to the upper chamber significantly impaired T cell migration toward CCL5 through narrow 5 µm pores (Fig. 4C), indicating a repulsive or inhibitory effect on chemokine-driven T cell migration. Interestingly, the addition of LCN2 to the lower chamber (acellular, CCL5-enriched) also resulted in a significantly reduced T cell influx, however, not to the same extent (Fig. 4C). This finding aligns with recent reports that LCN2 can modulate chemokine receptor signaling and limit T cell recruitment to inflamed tissues such as the TME (<u>Bossowski et al. 2026</u>) (85), potentially facilitating T cell egress from the spleen and thereby contributing to post-stroke immunodepression. Next, we examined the impact of LCN2 stimulation on monocytes isolated from the spleen. Again, pre-treatment with LCN2 significantly reduced their readiness to engage in LPS-induced up-regulation of *Il1b*, and (of borderline significance) *Tnf*, although no impact on *Il6* transcription was observed (Fig. 4D), corroborating the cytokine suppression in whole spleen homogenates post-tMCAO as suggested by the 40-plex cytokine array (Fig. 3B). This functional phenotype shift toward an anti-inflammatory or “tolerogenic” state is supported by studies showing that LCN2 suppresses classical macrophage activation via inhibition of the NF-κB-STAT3-axis, whilst promoting anti-inflammatory gene expression (<u>Guo et al. 2014</u>) (96). To further dissect the dose-dependent effects of LCN2, we analyzed splenocyte-derived monocytes (CD45^+^CD3^-^CD19^-^SSC^med^) either Ly6C^high^ vs. Ly6C^low^ exposed to vehicle or increasing concentrations of recombinant LCN2 protein. Subsequently, these were challenged with pHrodo-labeled *E*. *coli* bioparticles and assessed by flow cytometry (Supplementary Fig. 5A), thereby assessing changes to cell surface marker expression, phagocytic competence, and production of ROS required for adequate bacterial defense (<u>West et al. 2011</u>) (97). Indeed, LCN2 induced a reciprocal shift in Ly6C expression as the proportion of Ly6C^low^ (patrolling/repairing) monocytes increased (<u>Lessard et al. 2017</u>) (98). Conversely, Ly6C^high^ (inflammatory) monocytes decreased (<u>Lessard et al. 2017</u>) (98), with minimal changes in mean fluorescence intensity (MFI) being observed (Fig. 4E, Supplementary Fig. 5B). CD11b expression was selectively up-regulated in Ly6C^high^ monocytes (Fig. 4F, Supplementary Fig. 5C,D). In contrast, F4/80 expression decreased in this subset with higher LCN2 concentrations (Fig. 4G, Supplementary Fig. 5E,F), potentially consistent with a differentiation arrest in inflammatory monocytes (<u>Satoh et al. 2013</u>) (99). Functionally, LCN2 enhanced the phagocytic capacity of both monocyte subsets in a dose-dependent manner, as evidenced by increased uptake of *E*. *coli* bioparticles (Fig. 4H, Supplementary Fig. 5G). Notably, this increase in phagocytosis was uncoupled from ROS production. Whilst the percentage of ROS^+^ monocytes remained unchanged, the MFI of ROS was markedly reduced with higher LCN2 concentrations (Fig. 4I, Supplementary Fig. 5H,I). This uncoupling of phagocytosis from an oxidative burst response is a hallmark feature indicating a tolerogenic tone of LCN2-stimulated monocytes and may be a mechanism devised as a trade-off between pathogen clearance and minimizing tissue-damaging inflammation. Together, these findings suggest that LCN2 can orchestrate a coordinated reprogramming of both T cells and myeloid cells toward hyporesponsive, tolerogenic, and highly phagocytic phenotypes with an uncoupled oxidative burst response.

**Figure 4.**
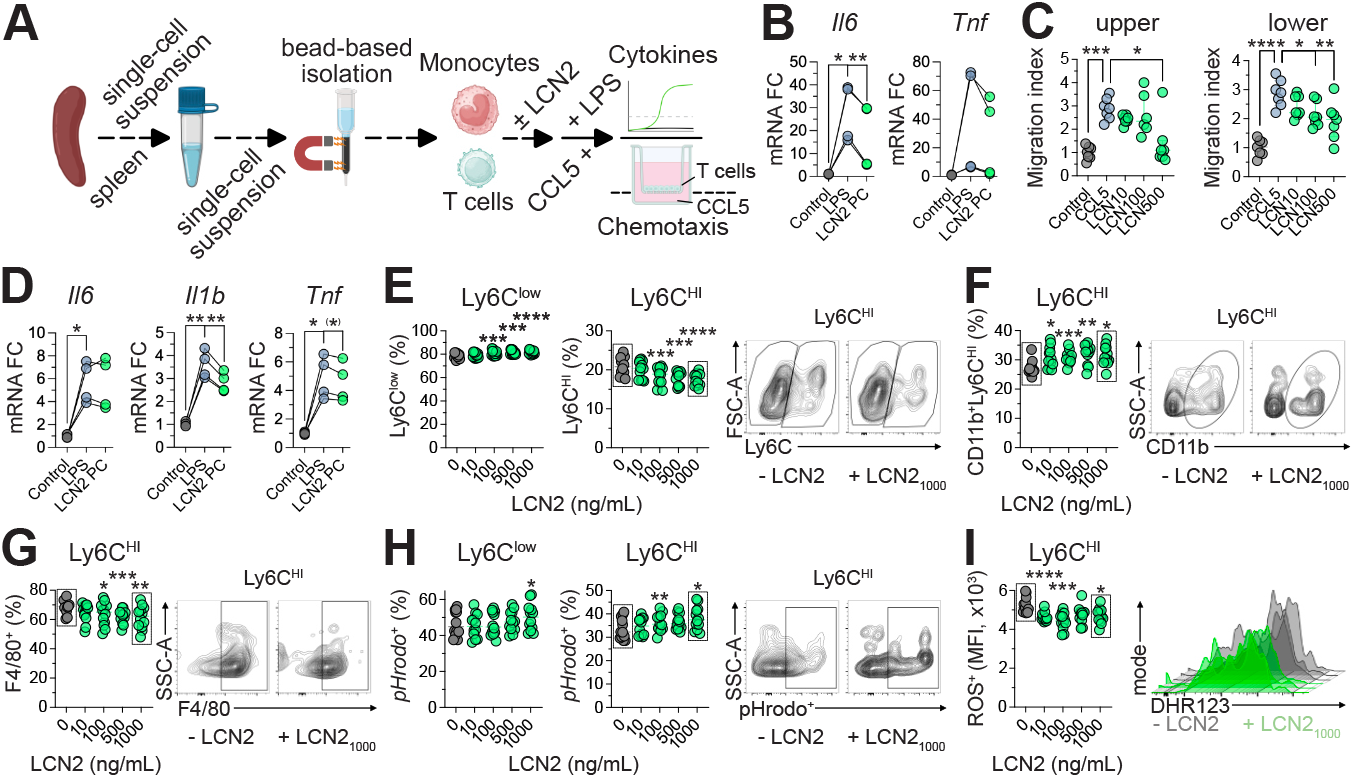
Lipocalin-2 induces hyporesponsive and phagocytosis-ROS-uncoupled phenotypes in splenic T cells and monocytes. (**A**) Experimental workflow: spleens from male, 10-12-week-old C57BL/6J mice subjected to tMCAO or sham surgery were processed into single-cell suspensions. T cells or monocytes were isolated magnetically and pre-conditioned with LCN2 or vehicle before LPS stimulation (100 ng/mL) to gauge LCN2-directed impact on LPS-induced proinflammatory cytokine production. For chemotaxis, 1×10^5^ T cells were tested in a 5-µm transwell assay toward CCL5 (200 ng/mL). (**B**) Fold-change expression of *Il6* or *Tnf* mRNA (ΔΔCT) normalized to *Gapdh* in T cells stimulated with LPS alone (blue) or after LCN2 preconditioning (green). Data (*n* = 4 per group) are symbols and lines and pooled from four independent experiments, paired *t* test; \**P* ≤ 0.05, \*\**P* ≤ 0.01. (**C**) Chemotaxis toward CCL5 with LCN2 placed in upper or lower chambers at 10-500 ng/mL. Data (*n* = 6-7 technical replicates, representative of two independent experiments) are median ± IQR. Kruskal-Wallis with Dunn’s test (upper) or one-way ANOVA with Dunnett’s test (lower); \**P* ≤ 0.05, \*\**P* ≤ 0.01, \*\*\**P* ≤ 0.001, \*\*\*\**P* ≤ 0.0001. (**D**) Fold-change *Il6*, *Il1b*, and *Tnf* mRNA in monocytes following LCN2 preconditioning and LPS stimulation. Data (*n* = 4 per group) are symbols and lines and pooled from four independent experiments, paired *t* test; ^(^*^)^*P* = 0.0513, \**P* ≤ 0.05, \*\**P* ≤ 0.01. (**E-I**) Flow cytometric analysis of monocytes treated with vehicle or increasing LCN2 doses. (**E**) Ly6C expression frequencies for Ly6C^low^ and Ly6C^HI^ monocytes. (**F**) CD11b expression on Ly6C^HI^ monocytes. (**G**) F4/80 expression on Ly6C^HI^ monocytes. (**H**) Phagocytic uptake (pHrodo^+^) in Ly6C^low^ and Ly6C^HI^ monocytes. (**I**) ROS production (DHR-123 MFI) during phagocytosis of *E*. *coli*. Representative plots/histograms are shown. Data (*n* = 12 per group, i.e., 6 technical replicates from two independent mice) are median ± IQR and pooled from two independent experiments; one-way ANOVA with Geisser-Greenhouse correction and Dunnett’s test; \**P* ≤ 0.05, \*\**P* ≤ 0.01, \*\*\**P* ≤ 0.001, \*\*\*\**P* ≤ 0.0001.

### Human ischemic stroke spleens show a conserved LCN2-expressing RPM-rich niche

Based on our murine findings that AIS induces LCN2 expression in splenic RPMs, we next asked whether this spatial organization is conserved in human ischemic stroke. Plasma samples from acute stroke patients showed elevated LCN2 levels at stroke onset and during follow-up compared with chronic post-stroke outpatient controls and healthy volunteers, consistent with previous reports identifying circulating LCN2 as a stroke-associated inflammatory biomarker (Fig. 5A,B) (<u>Hochmeister et al. 2016</u>; <u>Elneihoum et al. 1996</u>) (32,100). However, plasma measurements cannot identify the cellular source or tissue context of LCN2. We therefore analyzed post-mortem spleen sections from three patients with ischemic stroke with one non-stroke spleen included as a descriptive reference control. Immunofluorescence analysis revealed prominent LCN2 signal within CD68^+^ RPM-rich regions (Fig. 5C,D). Similar to the murine spleen after tMCAO (Fig. 3F,H), LCN2 was enriched in RPM-associated areas and was observed at close CD68^+^ macrophage-CD3^+^ T cell contact sites and CD68^+^ macrophage-CD68^+^ monocyte/macrophage contact sites (Fig. 5E,F). Quantitative image analyses showed consistent LCN2 enrichment in CD68^+^ RPM-rich regions featuring enriched areas for CD68^+^ monocytes and/or macrophages, LCN2^+^ area, and CD68^+^LCN2^+^ area across all three human ischemic stroke spleens (Fig. 5G, Supplementary Figure 6). Although the limited sample size and observational nature of post-mortem human tissue precludes causal inference, these data provide direct spatial support that the splenic LCN2-expressing RPM niche identified in experimental stroke is also present in human AIS.

**Figure 5.**
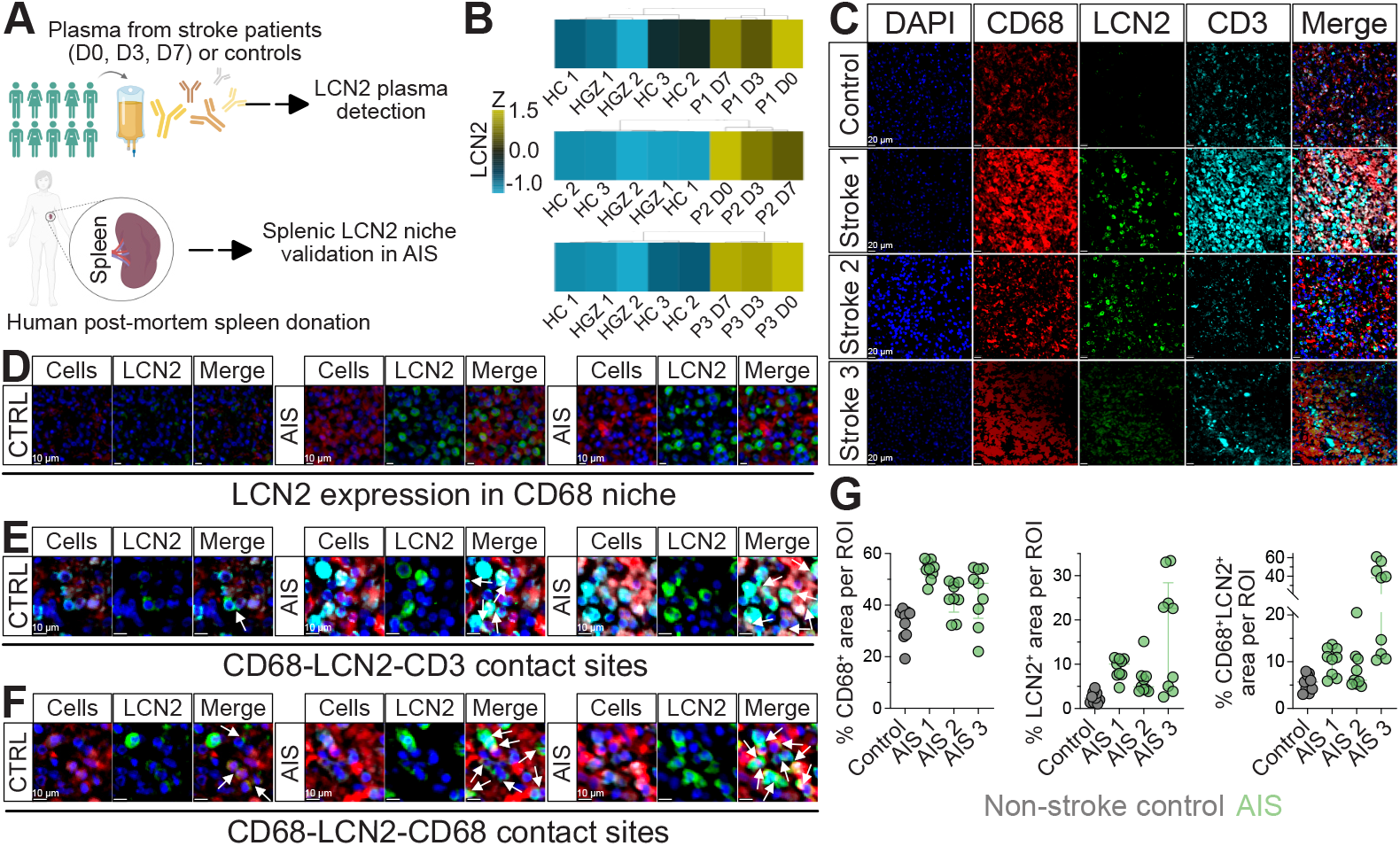
Lipocalin-2 expression by CD68^+^ red pulp macrophages is conserved in human ischemic stroke spleens. (**A**) Schematic of the study characteristics. Plasma from stroke patients at the day of stroke onset (D0) or after 3 (D3) or 7 days (D7) or patients months after stroke in the outpatients’ department (HGZ) or healthy volunteers (HC) was collected. Plasma samples were analyzed for Lipocalin-2 (LCN2) expression using an IgG-based detection system by OmicsArray^TM^ antigen array. Post-mortem spleen sections from three patients with AIS and one non-stroke reference spleen were analyzed by immunofluorescence. (**B**) Clustering of soluble LCN2 expression in plasma of patients 1-3 (P1-3) across the different timepoints following stroke onset as compared to HGZ or HCs showing highest LCN2 expression at onset, lower expression at D3 with regained stronger expression at D7, from three independent experiments. (**C**) Representative immunofluorescence of human spleen sections as indicated in (**A**). Individual channels for DAPI, CD68, LCN2, CD3, and their merged composite (Merge) are shown. Scale bar = 20 µm. (**D-F**) Detailed images from non-stroke reference control (CTRL) or postmortem spleens from patients with acute ischemic stroke (AIS). Scale bar = 10 µm. (**D**) Representative high-resolution images showing LCN2 enrichment in the CD68^+^ RPM-rich niche upon AIS, indicated by white arrowheads. (**E**) Representative high-resolution images showing LCN2-enriched CD68^+^ macrophage-associated contact sites with CD3^+^ T cells in AIS, indicated by white arrowheads. (**F**) Representative high-resolution images showing LCN2-enriched CD68^+^ macrophage-associated contact sites with CD68^+^ monocyte/macrophage cells, indicated by white arrowheads. (**G**) Patient-level quantification of CD68^+^, LCN2^+^, or CD68^+^/LCN2^+^ area with-in annotated red pulp regions of interest (ROI). Data (*n* = 9 ROIs per group) are median ± IQR. Each ROI input is shown in Supplementary Figure 6. The non-stroke spleen is shown as a descriptive reference control only; no formal statistical inference was performed for stroke versus non-stroke spleen comparisons.

### LCN2 induces tolerogenic and phagocytosis-ROS-uncoupled reprogramming of human monocytes

Having established that LCN2-expressing CD68^+^ RPM-rich niches are present in human ischemic stroke spleens, we next tested whether LCN2 is sufficient to reprogram human monocytes in a reductionist *ex vivo* system. To directly assess the functional impact of LCN2, untouched monocytes were isolated from healthy donor PBMCs using negative selection (Fig. 6A). These monocytes were then exposed *in vitro* to increasing concentrations of recombinant LCN2 protein, modeling the acute-phase exposure encountered in the post-stroke spleen milieu (Fig. 5D,F). Flow cytometric profiling revealed that LCN2 did not alter the relative proportions of classical (CD14^high^) and intermediate (CD14^dim^) monocyte subsets (Fig. 6B, Supplementary Fig. 7A), but induced a striking, dose-dependent modulation of their phenotypic and functional properties. Specifically, CD14 MFI increased on intermediate monocytes and decreased on classical monocytes (Fig. 6C), suggesting an altered TLR4 co-receptor availability and potentially a dampened inflammatory readiness in the latter subset (<u>Nielsen et al. 2020</u>) (101). MFI of CD11b surface expression was dramatically reduced in both populations, despite unchanged percentages of CD11b^+^ monocytes (Fig. 6D, Supplementary Fig. 7B), indicating an impaired adhesion and migratory potential, a feature associated with monocyte retention and reduced tissue infiltration in immunosuppressed states (<u>Nielsen et al. 2020</u>) (101). A hallmark of post-stroke immunodepression is the down-regulation of HLA-DR, a key antigen-presenting molecule to CD4^+^ T helper cells expressed by circulating monocytes and tissue-resident macrophages (<u>Kaito et al. 2013</u>; <u>Urra et al. 2009</u>) (102,103). Consistent with this, LCN2 exposure led to a significant, dose-dependent reduction in both the percentage and MFI of HLA-DR on classical and intermediate monocytes (Fig. 6E, Supplementary Fig. 7C). This loss of antigen presentation capacity is clinically linked to impaired T cell activation and increased infection risk in stroke patients (<u>Kaito et al. 2013</u>; <u>Urra et al. 2009</u>) (102,103). Similarly, CD80, a critical co-stimulatory molecule to mount effector T cell responses (<u>Odobasic et al. 2008</u>; <u>Narni-Mancinelli et al. 2007</u>) (104,105), was reduced in both monocyte subsets with increasing LCN2 concentrations (Fig. 6F), further supporting a shift towards a hyporesponsive, tolerogenic phenotype (<u>Li et al. 2024</u>) (106). LCN2 also modulated CD206, a macrophage marker associated with impaired tissue functions including in ischemic stroke (<u>Wang et al. 2025</u>; <u>Nawaz et al. 2022</u>; <u>Drieu et al. 2020</u>) (107–109). CD206 expression was significantly reduced on classical monocytes at higher LCN2 concentrations but remained unchanged with lower LCN2 levels or on intermediate monocytes (Fig. 6G, Supplementary Fig. 7D). Functionally, LCN2 dramatically enhanced the phagocytic capacity of both monocyte subsets in a dose-dependent manner, as measured by the uptake of pHrodo-labeled *E*. *coli* bioparticles (Fig. 6H, Supplementary Fig. 7E). Importantly, this increased phagocytic competence was again uncoupled from ROS production consistent with our observations in monocytes isolated from C57BL/6J mice (Fig. 4H,I). Indeed, whilst the proportion of ROS^+^ monocytes was unchanged, ROS MFI decreased in classical monocytes with LCN2 stimulation, although at higher concentrations only with borderline statistical significance (Fig. 6I, Supplementary Fig. 7F). This uncoupling of ROS, i.e., phagocytic uptake with a suppressed oxidative burst response, is a defining feature of tolerogenic myeloid cells, aimed to enable pathogen clearance whilst minimizing collateral tissue damage (<u>Vassallo et al. 2019</u>) (110). Together, these findings have potential clinical relevance because monocyte dysfunction, characterized by HLA-DR down-regulation, impaired antigen presentation, and reduced T cell activation, represents a central mechanism underlying post-stroke immunodepression and increased infection risk in humans (<u>Westendorp et al. 2011</u>) (10). The ability of recombinant LCN2 to induce antigen-presentation loss, co-stimulatory marker reduction and phagocytosis-ROS uncoupling in a cell-intrinsic manner in human monocytes supports a model in which LCN2 contributes to systemic immune conditioning after AIS and may link acute-phase responses to post-stroke monocyte dysfunction.

**Figure 6.**
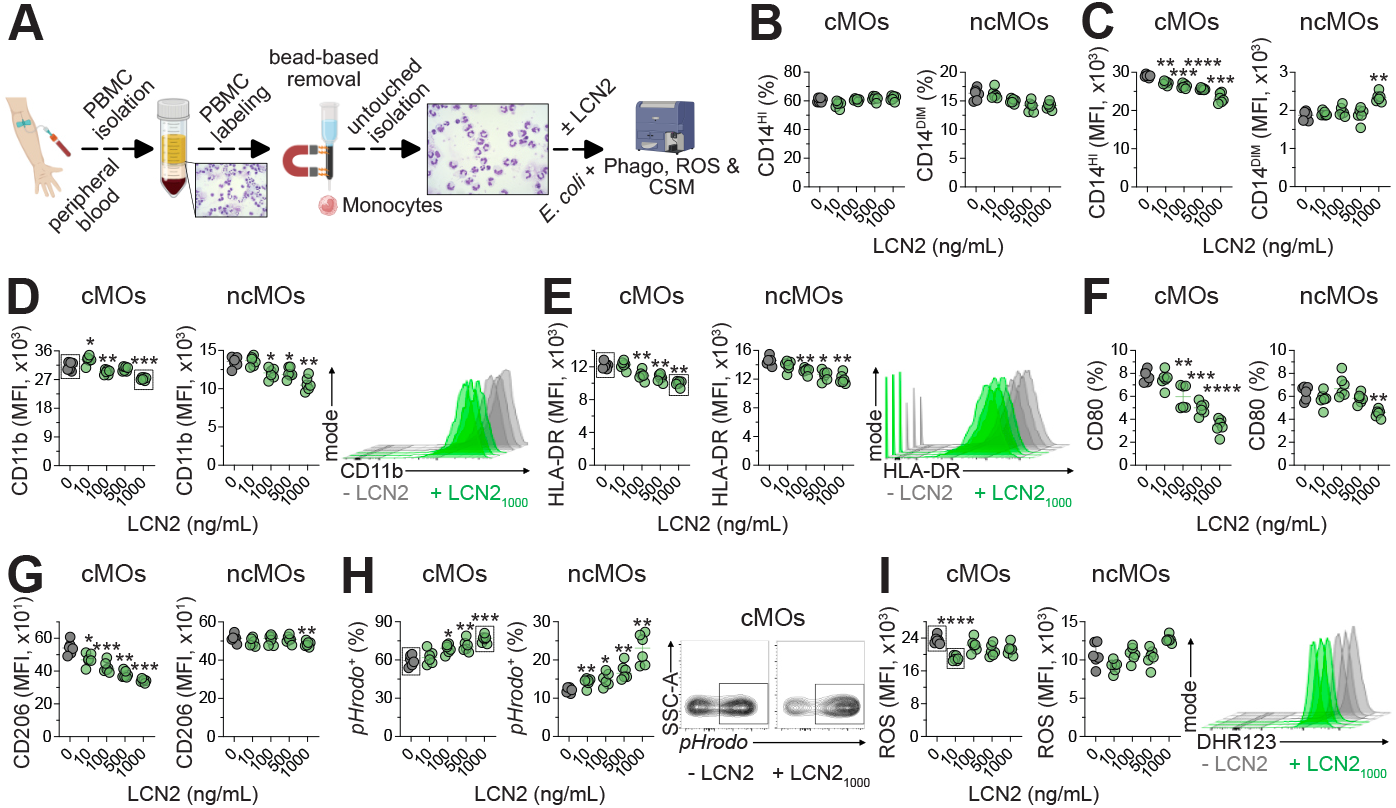
Lipocalin-2 induces tolerogenic and phagocytosis-ROS-uncoupled reprogramming of human monocytes. (**A**) Experimental workflow: peripheral blood from healthy donors was processed by Ficoll-based density gradient centrifugation to isolate PBMCs, followed by untouched monocyte enrichment using a negative-selection kit. Cytospins before/after isolation confirmed purity. Monocytes were pre-treated with vehicle or LCN2 (escalating concentrations) and challenged with pHrodo-labeled *E*. *coli*. After 30 minutes, phagocytosis was stopped, and ROS (DHR-123), phagocytic uptake, and expression of cell-surface markers (CSM) were assessed by flow cytometry. (**B**) Frequencies of CD14^DIM^ or CD14^HI^ (intermediate (ncMOs) or classical monocytes (cMOs), respectively) after LCN2 stimulation for 4 hours. (**C**) CD14 MFI in cMOs and ncMOs. (**D**) CD11b MFI in both subsets with representative histograms. (**E**) HLA-DR MFI in cMOs and ncMOs with representative histograms. (**F**) Percentage of CD80 in cMOs or ncMOs. (**G**) CD206 MFI in both subsets. (**H**) Phagocytic uptake (pHrodo^+^) in cMOs and ncMOs with representative contour plots. (**I**) ROS MFI during *E*. *coli* exposure in cMOs and ncMOs with representative histograms. (**B-I**) Data (*n* = 6 technical replicates per group) are median ± IQR and representative of two independent experiments (biological donors); Friedman with Dunn’s test (cMOs in **B**) or one-way ANOVA with Geisser-Greenhouse correction and Dunnett’s test; \**P* ≤ 0.05, \*\**P* ≤ 0.01, \*\*\**P* ≤ 0.001, \*\*\*\**P* ≤ 0.0001.

## Discussion

Here, we identify splenic LCN2-expressing RPMs as a conserved cellular hub of post-stroke peripheral immune cell reprogramming. In murine tMCAO, *Lcn2* was among the most strongly induced splenic transcripts and showed biphasic regulation during the acute and delayed post-stroke phases. Spatial analysis localized LCN2 predominantly to F4/80^+^ RPMs, where it accumulated at RPM-associated T cell contact sites. Importantly, analysis of post-mortem spleens from three patients with ischemic stroke validated the presence of LCN2-expressing CD68^+^ RPM-rich regions in human AIS. Together with elevated circulating LCN2 in stroke plasma and functional reprogramming of murine and human immune cells by recombinant LCN2, these data position LCN2 as a translationally conserved candidate mediator linking cerebral ischemia to systemic immune cell dysfunction. These observations extend previous work implicating the spleen as a central modulator of post-stroke immunity, where structural and functional changes, including atrophy and immune cell depletion have been described as consequences of neuro-endocrine activation and sympathetic signaling following ischemic stroke (<u>Offner et al. 2006a</u>; <u>Mracsko et al. 2014</u>; <u>Mcculloch et al. 2017</u>) (18,21,22). Our data reveal that LCN2-mediated immune cell conditioning is not restricted to the CNS but represents a systemic phenomenon with direct consequences for peripheral immunity.

Mechanistically, we observed that LCN2 reduced proinflammatory cytokines in T cells and monocytes, impaired T cell diapedesis, and uncoupled phagocytic uptake from oxidative burst responses in monocytes following exposure to *E*. *coli*, a gram-negative rod often responsible for post-stroke infections (<u>Westendorp et al. 2011</u>; <u>Elkind et al. 2020</u>) (10,111). These findings are consistent with prior studies demonstrating that LCN2 can induce immune tolerance in both innate and adaptive immune compartments, potentially through modulation of iron homeostasis, inflammatory signaling, and metabolic reprogramming (<u>Devireddy et al. 2005</u>; <u>Watzenboeck et al. 2021</u>; <u>Marques et al. 2023</u>) (33,39,112). The ability of LCN2 to uncouple phagocytosis from ROS production in monocytes suggests a mechanism by which bacterial-directed phagocytosis can occur without causing excessive tissue-damaging inflammation, a hallmark of tolerogenic myeloid responses (<u>Restaino et al. 2017</u>; <u>Hanlon et al. 2023</u>) (113,114), however, as a potential trade-off for antimicrobial activity and intracellular bacterial degradation (<u>West et al. 2011</u>; <u>Cavinato et al. 2020</u>) (97,115). Moreover, our data indicate that LCN2 impairs T cell migration toward chemokines such as CCL5, thereby limiting the recruitment of effector T cells to sites of inflammation. This is in line with recent findings that LCN2 can modulate chemokine availability and thus immune cell recruitment (<u>Borkham-Kamphorst et al. 2021</u>) (116).

The infection-related implications of splenic LCN2 induction require careful interpretation. Our murine and human data show that recombinant LCN2 increases gram-negative bacterial particle uptake by monocytes whilst reducing the phagocytosis-induced oxidative burst response (<u>Yi et al. 2022</u>; <u>Herb et Schramm 2021</u>) (117,118), suggesting an uncoupling of phagocytosis from oxidative antimicrobial effector function. This phenotype provides a plausible mechanistic link between LCN2-driven post-stroke immune conditioning and infection susceptibility, particularly because impaired monocyte antigen presentation and altered innate effector functions are established features of post-stroke immunodepression. However, LCN2 also has pathogen-restrictive functions through siderophore sequestration and has been reported to protect against selected bacterial infections, including *E*. *coli* and *K*. *pneumoniae* (<u>Zhang et al. 2023</u>; <u>Wu et al. 2010</u>; <u>Cramer et al. 2017</u>) (44–46). Conversely, LCN2 can deactivate macrophages and worsen *S*. *pneumoniae*-induced pneumonia outcomes, and can modulate dendritic cell activity and antiviral immunity to influenza in a context-dependent manner (<u>Warszawska et al. 2013</u>; <u>Watzenboeck et al. 2021</u>) (38,39). Thus, the net effect of LCN2 on infection susceptibility is likely pathogen-, compartment-, timing– and receptor-dependent.

Importantly, the human spleen data strengthen the translational relevance of this axis. While plasma LCN2 measurements indicate systemic induction after stroke, they cannot identify the cellular source or spatial immune context of LCN2. Detection of LCN2 in CD68^+^ RPM-rich regions of human ischemic stroke spleens provides direct anatomical evidence that the macrophage-associated LCN2 niche observed in tMCAO is also present in human AIS. Together with the ability of recombinant LCN2 to reduce HLA-DR, CD80, CD206, and CD11b expression in human monocytes, these data support a model in which LCN2 contributes to post-stroke peripheral immune cell conditioning. From a mechanistic standpoint, our study extends the current understanding of CIDS, a phenomenon characterized by lymphocytopenia, impaired antigen presentation, and increased infection susceptibility (<u>Meisel et al. 2005</u>; <u>Prass et al. 2003</u>; <u>Urra et al. 2009</u>) (8,9,103). LCN2 emerges as a pivotal factor in driving CIDS given its magnitude and timing of upregulation in RPMs, and contribution to cell-to-cell contact sites, thereby adding to its functions as an acute-phase protein or regulator of iron metabolism (<u>Yang et al. 2002</u>; <u>Ferreira et al. 2015</u>) (35,119).

Together, our observations nominate LCN2 signaling as a candidate therapeutic axis for future studies aimed at mitigating CIDS and infection-related morbidity (<u>Zhang et al. 2023</u>; <u>Wu et al. 2010</u>; <u>Cramer et al. 2017</u>) (44–46). While our study provides comprehensive insights into the temporal and mechanistic landscape of LCN2-mediated splenic immune reprogramming after stroke, several important limitations must be acknowledged.

First, although the use of post-mortem human ischemic stroke spleens provides important translational support, the sample size remains limited and the observational nature of these tissues preclude causal inference. Post-mortem interval, stroke severity, comorbidities, infection status, and treatment history may influence splenic immune architecture and LCN2 expression. Therefore, the human spleen data should be interpreted as spatial support of the LCN2-expressing RPM niche, rather than proof of causality.

Second, our functional human assays used monocytes from healthy donors exposed to recombinant LCN2. This controlled approach demonstrates LCN2 sufficiency but does not fully recapitulate the complex plasma and tissue milieu of AIS patients. Longitudinal analyses of patient-derived monocytes, infection outcomes, and matched circulating LCN2 levels will be required to define whether LCN2-dependent monocyte and T cell remodeling predict post-stroke infection susceptibility.

Third, our phagocytosis assays used pHrodo-labeled *E*. *coli* bioparticles and DHR-123-based ROS detection as functional surrogates of bacterial uptake and oxidative antimicrobial competence. These assays demonstrate that LCN2 enhances phagocytic uptake while reducing ROS intensity, but they do not directly measure intracellular ability for microbial killing. Live-pathogen assays using CFU readouts for bacteria and fungi, and replication-based readouts for viruses, will be required to determine whether this phenotype causally alters infection susceptibility.

Fourth, while our data implicate LCN2 as a candidate mediator of splenic immune reprogramming, the receptor-specific contributions of 24p3R and LRP2 remain unresolved. Future studies using receptor-specific genetic perturbation, validated blocking approaches, and cell-type-specific *in vivo* models will be required to define whether LCN2 signaling is necessary for post-stroke immune dysfunction.

Fifth, indiscriminate LCN2 blockade may be undesirable because LCN2 contributes to antimicrobial host defense through pathogen– and context-dependent mechanisms (<u>Nairz et al. 2015</u>; <u>Chan et al. 2009</u>; <u>Moschen et al. 2016</u>) (41–43). Therapeutic approaches should therefore prioritize time-restricted, cell-type-specific or receptor-selective modulation of LCN2 signaling rather than global depletion.

Finally, our experimental paradigm primarily utilized the tMCAO-model in young, healthy C57BL/6J mice. Although tMCAO is a well-established model for ischemic stroke, it does not fully recapitulate the heterogeneity of human stroke, which encompasses ischemic and hemorrhagic subtypes, variable reperfusion dynamics, and a broad spectrum of comorbidities such as hypertension, diabetes, and advanced age. These factors are known to profoundly influence both stroke outcomes and immune responses, with aged and comorbid animals exhibiting more severe immunosuppression, altered splenic dynamics, and increased susceptibility to infection compared to young, healthy counterparts. Future studies should incorporate aged and comorbid animal models, as well as alternative stroke paradigms, including ischemic and hemorrhagic models to enhance translational relevance and to capture the diversity of clinical stroke presentations.

## Conclusions

AIS induces a profound remodeling of peripheral immunity, and our data identify LCN2-expressing RPMs as a conserved splenic immunoregulatory niche in mice and humans. In murine tMCAO, splenic *Lcn2* was rapidly induced, localized predominantly to RPMs and accumulated at RPM-associated T cell contact sites. Human ischemic stroke spleens support the presence of LCN2-expressing CD68^+^ RPM-rich regions, supporting translational conservation of this axis. Functionally, recombinant LCN2 reprogrammed murine T cells and monocytes and human monocytes in a cell-intrinsic manner towards hyporesponsive, tolerogenic, and highly phagocytic states, characterized by suppressed proinflammatory cytokine production, impaired chemotactic responses, reduced antigen-presentation and co-stimulatory markers, and uncoupling of phagocytosis from ROS production.

Together, these findings establish LCN2 as a candidate mechanistic link between ischemia-induced CNS injury and systemic immune dysfunction, thus positioning LCN2-expressing RPMs as a potential therapeutic axis in CIDS. Because LCN2 also contributes to antimicrobial host defense through pathogen– and context-dependent mechanisms, future therapeutic approaches should avoid indiscriminate LCN2 depletion. Instead, studies should define whether time-restricted, cell-type-specific or receptor-selective modulation of LCN2 signaling through 24p3R or LRP2 can preserve antimicrobial protection while preventing LCN2-driven monocyte and T cell hyporesponsiveness.

## Supplementary Figures

**Supplementary Figure 1.**
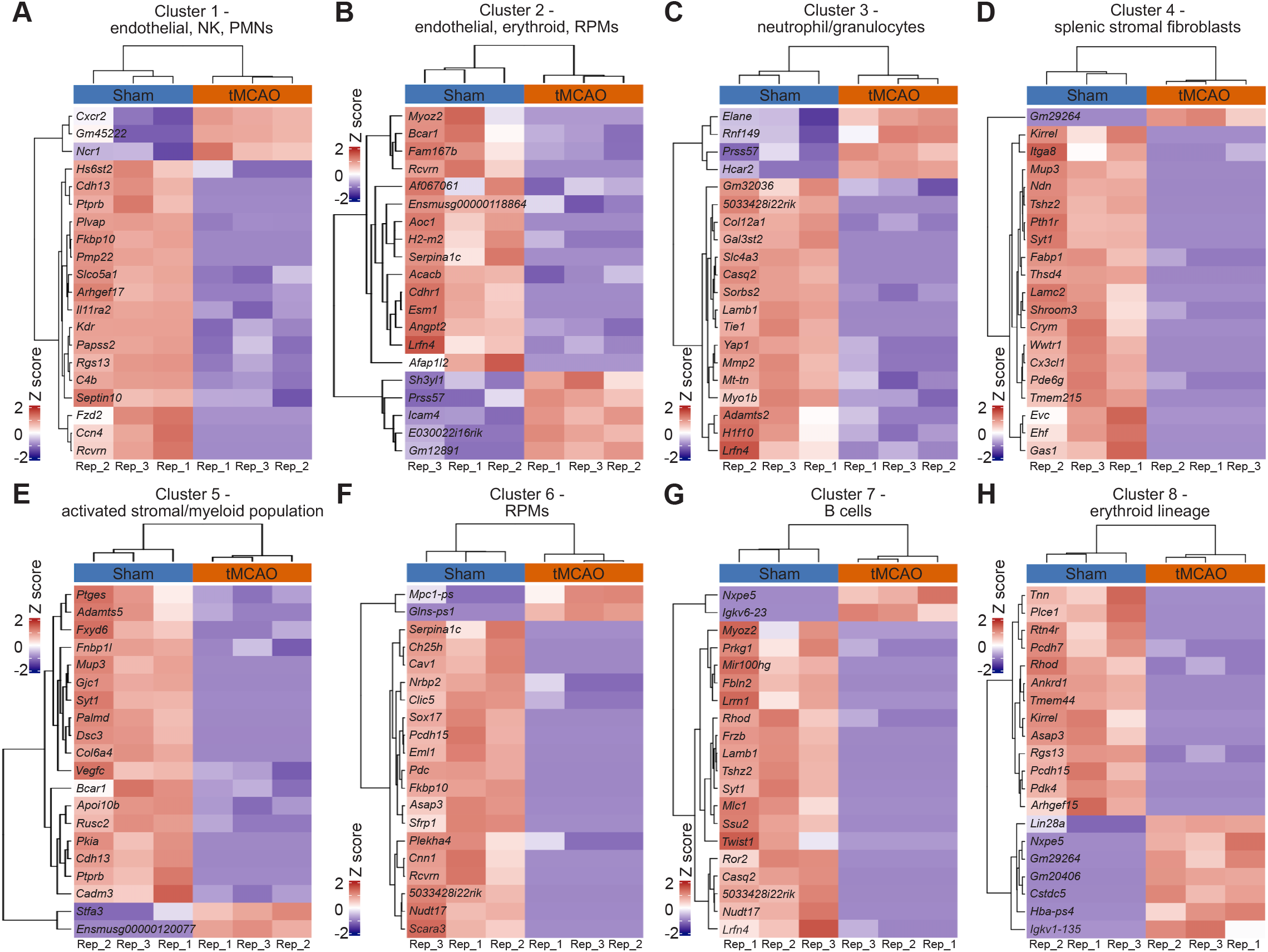
– related to Figure 1. Top marker genes by k-means clustering of 3’-MACE-Seq data from sham– and tMCAO-treated mice. (**A-H**) Heatmaps display the Z scores of the top 20 marker genes across experimental conditions (sham-operation and tMCAO) and biological replicates. (**A**) Canonical markers identify cluster 1 to mark sinusoidal endothelial cells (*Kdr*, *Plvap*), NK cells (*Ncr1*), and neutrophils (*Cxcr2*), indicating a mixed vascular and innate immune population. (**B**) Canonical markers identify cluster 2 to characterize transcripts pertaining to the erythroid lineage (*Icam4*), endothelial cells (*Esm1*, *Angpt2*), and with red pulp macrophage (RPM)-association (*H2-m2*). Also, it contains non-immune markers such as *Myoz2* and *Rcvrn*. (**C**) Canonical markers identify cluster 3 to robustly define neutrophil/granulocyte markers (*Elane*, *Prss57*, *Hcar2*). (**D**) Canonical markers identify cluster 4 to be enriched for transcripts associated with stromal fibroblasts and perivascular markers (*Fabp1*, *Cx3cl1*, *Kirrel*, *Itga8*, *Gas1*). (**E**) Canonical markers identify cluster 5 to display a profile of activated stromal/endothelial and myeloid genes (*Ptges*, *Adamts5*), but lacking canonical granulocyte markers. (**F**) Canonical markers identify cluster 6 to incorporate RPM-associated transcripts (*Ch25h*, *Serpina1c*, *Cav1*, *Sox17*). (**G**) Canonical markers identify cluster 7 to mark B cells as indicated by B cell-specific immunoglobulin genes (*Igkv6-23*). (**H**) Canonical markers identify cluster 8 to characterize erythroid progenitors (*Hba-ps4*, *Lin28a*, *Rhod*).

**Supplementary Figure 2.**
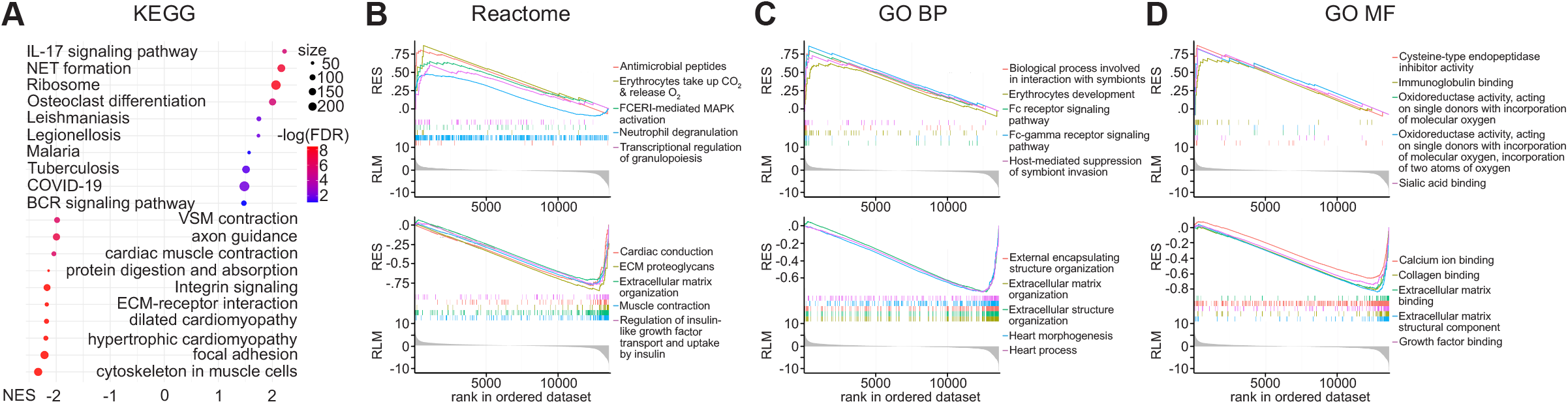
– related to Figure 1. Multi-database GSEA in the spleen following acute ischemic stroke. (**A**) Bubble plot of significantly enriched KEGG pathways comparing 24 hour tMCAO versus sham-operated spleens. The x-axis displays the normalized enrichment score (NES), and pathway names are indicated. The bubble size denotes the gene size within each pathway and the color intensity reflects the FDR. (**B-D**) Enrichment plots for the top 5 up-regulated (top) and top 5 down-regulated pathways (bottom) using the Reactome database (**B**), the database for Gene Ontology Biological Processes (GO:BP) (**C**), or the database for Gene Ontology Molecular Function (GO:MF) (**D**). Each plot displays the running enrichment score (RES) curve, the ranked position of individual pathway genes in the ordered gene list, and a heatmap of normalized expression values (RLM).

**Supplementary Figure 3.**
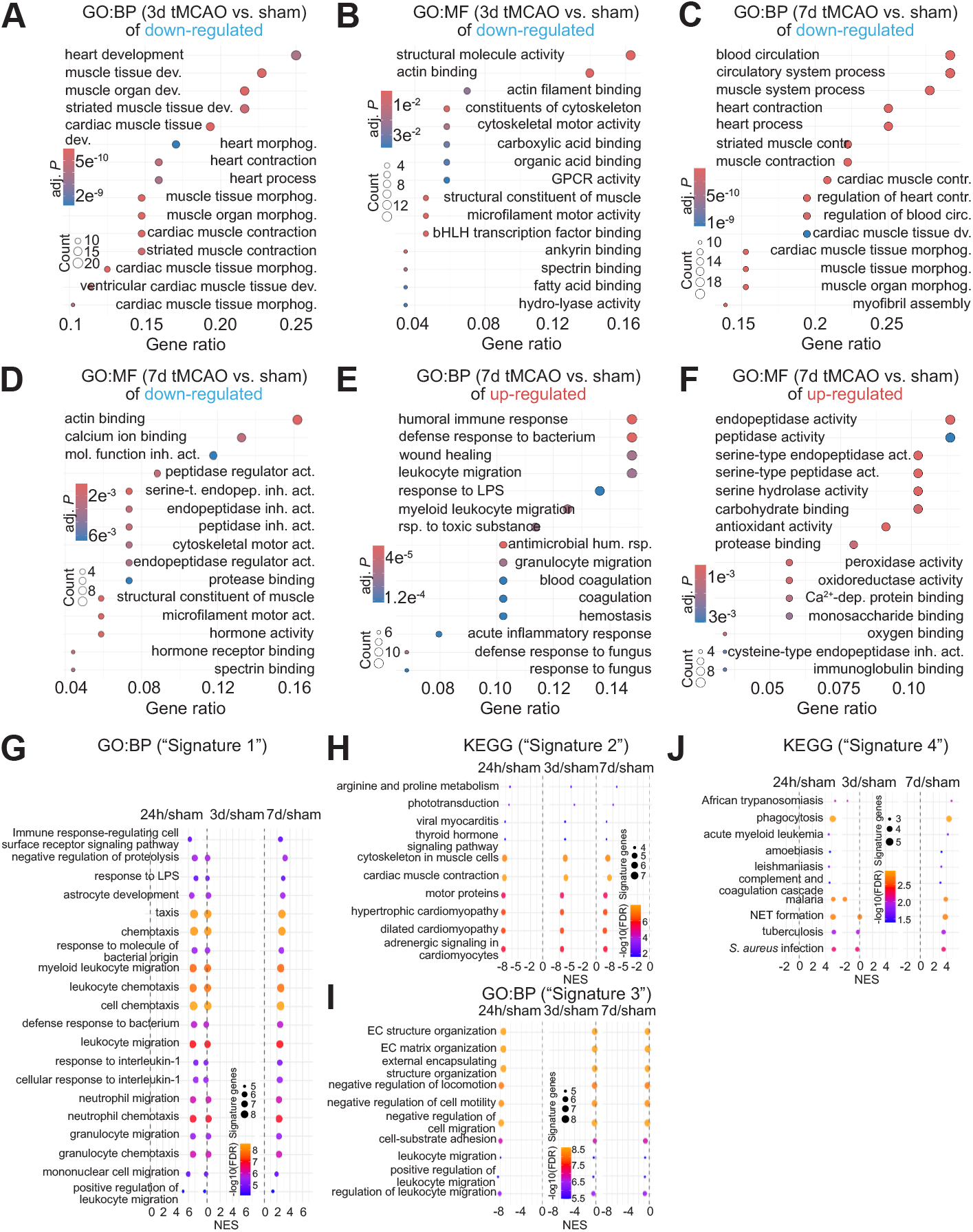
– related to Figure 2. Enrichment analyses show differential regulation of immune pathways and structural remodeling in the spleen following acute ischemic stroke. (**A-B**) Bubble plot of significantly depleted GO:BP (**A**) or GO:MF (**B**) pathways comparing spleen sequencing results 3 days after tMCAO (3d tMCAO) versus sham-operated animals (sham). (**C-D**) Bubble plot of significantly depleted GO:BP (**C**) or GO:MF (**D**) pathways comparing spleen sequencing results 7 days after tMCAO (7d tMCAO) versus sham-operated animals (sham). (**E-F**) Bubble plot of significantly enriched GO:BP (**E**) or GO:MF (**F**) pathways comparing spleen sequencing results 7 days after tMCAO (7d tMCAO) versus sham-operated animals (sham). (**A-F**) The x-axis displays the gene ratio (proportion of genes in the term amongst all genes), and pathway names are indicated. The bubble size denotes the gene count within each pathway and the color intensity reflects the adjusted *P* value (adj. *P*). Abbreviations: dev./dv. Development; morphog., morphogenesis; contr., contraction; circ., circulation; act., activity; inh., inhibitor; mol., molecular; rsp., response; hum., humoral. (**G**) Top GO:BP terms associated with “Signature 1” following GSEA, showing the numerical enrichment score (NES) on the x-axis and the size of identified genes within each pathway and the false discovery rate (FDR). (**H**) Analysis of “Signature 2” using KEGG, showing NES on the x-axis and the size of identified genes within each pathway and the FDR. (**I**) Same as (**G**) using “Signature 3”. (**J**) Same as (**H**) using “Signature 4”.

**Supplementary Figure 4.**
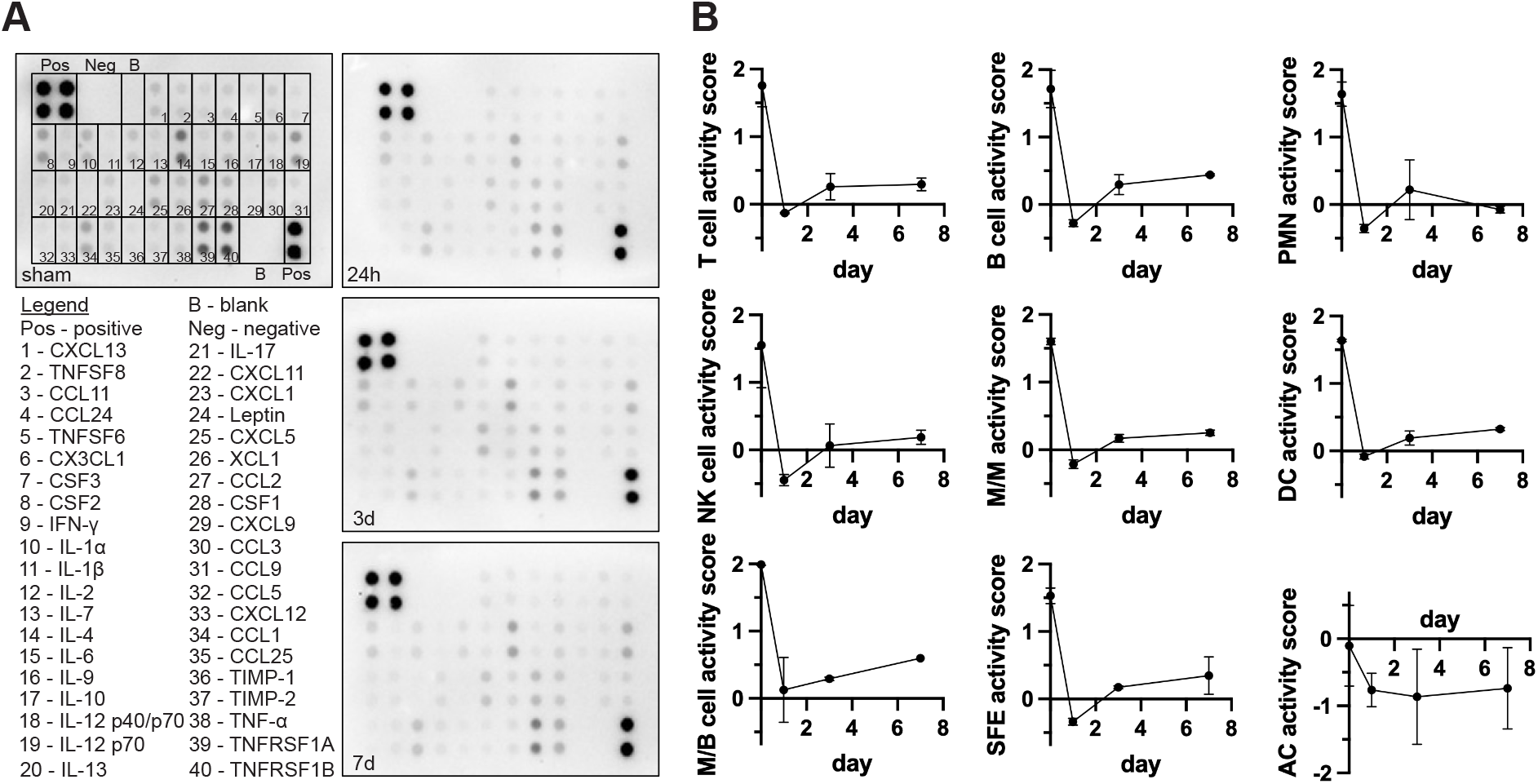
– related to Figure 3. Lipocalin-2 contributes to multicellular effector cell activity in the spleen after ischemic stroke. (**A**) Representative membranes from the performed Cytokine array done on 100 µg of spleen lysates isolated from sham-operated or 24 hours, 3 days, or 7 days following tMCAO-treatment in C57BL/6J mice. The legend illustrates the 40 cytokines detected in the assay as exemplary marked on the sham membrane. (**B**) The cytokines associated with T cells (IFNγ, IL-2, IL-4, IL-9, IL-10, IL-13, IL-17, FASL, CD30L, CCL5, XCL1, TCA-3, TNFα, CSF2), B cells (IL-10, MIP-1α, CCL5), polymorphonuclear neutrophils (PMNs; IL-1β, KC, LIX), NK cells (IFNγ, FASL, XCL1, CCL5), macrophages/monocytes (M/M; IL-1α, IL-1β, IL-6, IL-10, TNFα, MCP-1, MIP-1α, MIP-1γ, KC, LIX, MIG, I-TAC, CSF1, TIMP-1, TIMP-2, sTNF RI, sTNF RII, CSF2), dendritic cells (DCs; IL-12 p70, IL-12 p40/p70, IL-1α, IL-1β, IL-6, IL-10, MCP-1, MIP-1α, MIP-1γ, MIG, I-TAC, TCA-3, TECK, CD30L, TNFα, CSF2), mast cells/basophils (M/B; IL-4, IL-9, IL-13), stromal cells/FRCs/endothelial cells (SFE; BLC, CCL11, CCL24, CX3CL1, CSF1, CSF3, IL-7, SDF-1, TECK, TIMP-1, TIMP-2, sTNF RI, sTNF RII, leptin), or adipocytes (ACs; leptin) were combined towards a cell-specific activity score to kinetically map effector cell activity over the course of days following acute ischemic stroke. Data (*n* = 2 per group; each *n* represents four biologically distinct mice) are mean ± SD and representative of two independent experiments. Statistical comparisons were not performed.

**Supplementary Figure 5.**
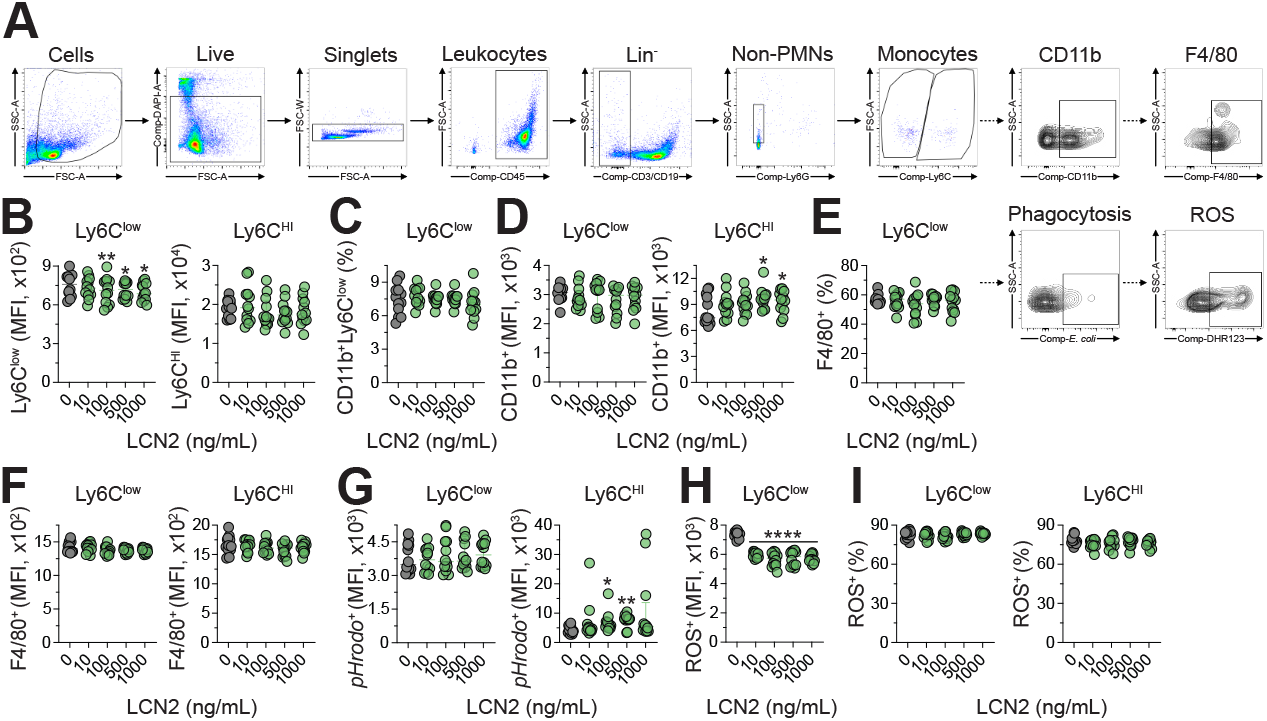
– related to Figure 4. Exposure to recombinant lipocalin-2 is linked to immunomodulation of monocytes in the spleen. (**A**) Gating strategy for flow cytometry to identify cells by FSC-A and SSC-A to exclude debris followed by identification of live cells (DAPI negative) and selection of singlets (FSC-A vs. FSC-W). Next, leukocytes were identified by CD45 positivity followed by selection of CD3/CD19 lineage-negative cells, and subsequent identification of non-PMN (Ly6G-negative), SSC-A-medium monocytes. These were used for characterization of CD11b and F4/80 expression or ability to phagocytose *E*. *coli* bioparticles or produce reactive oxygen species (ROS). (**B**) Mean fluorescent intensity (MFI) of Ly6C^low^– (left) or Ly6C^HI^-positive monocytes (right) within C57BL/6J splenocytes either vehicle– or LCN2-stimulated at the indicated concentrations for 4 hours prior to flow cytometric analysis. (**C**) Same as (**B**) showing CD11b expression (in %) on Ly6C^low^ monocytes. (**D**) Same as (**B**) but showing MFI of CD11b. (**E**) Same as (**B**) showing F4/80 expression (in %) on Ly6C^low^ monocytes. (**F**) Same as (**B**) showing F4/80 expression (in MFI) on Ly6C^low^ (left) or Ly6C^HI^ monocytes (right). (**G**) Phagocytic competence as indicated by MFI for pHrodo^+^ Ly6C^low^ (left) or Ly6C^HI^ monocytes following a 30-minute exposure to pHrodo-labeled *E*. *coli* bioparticles (25 µg/mL). (**H**) MFI of ROS as measured by DHR-123-FITC during exposure to *E*. *coli* bioparticles in Ly6C^low^ monocytes. (**I**) Expression of ROS (in %) as measured by DHR-123-FITC during exposure to *E*. *coli* bioparticles in Ly6C^low^ (left) or Ly6C^HI^ monocytes (right). (**B-I**) Data (*n* = 12 per group, i.e., 6 technical replicates from two independent mice) are median ± IQR and pooled from two independent experiments. Statistical comparisons were performed with a one-way ANOVA, with Tukey’s multiple comparison test for parametric data. \**P* ≤ 0.05, \*\**P* ≤ 0.01, \*\*\**P* ≤ 0.001, \*\*\*\**P* ≤ 0.0001.

**Supplementary Figure 6.**
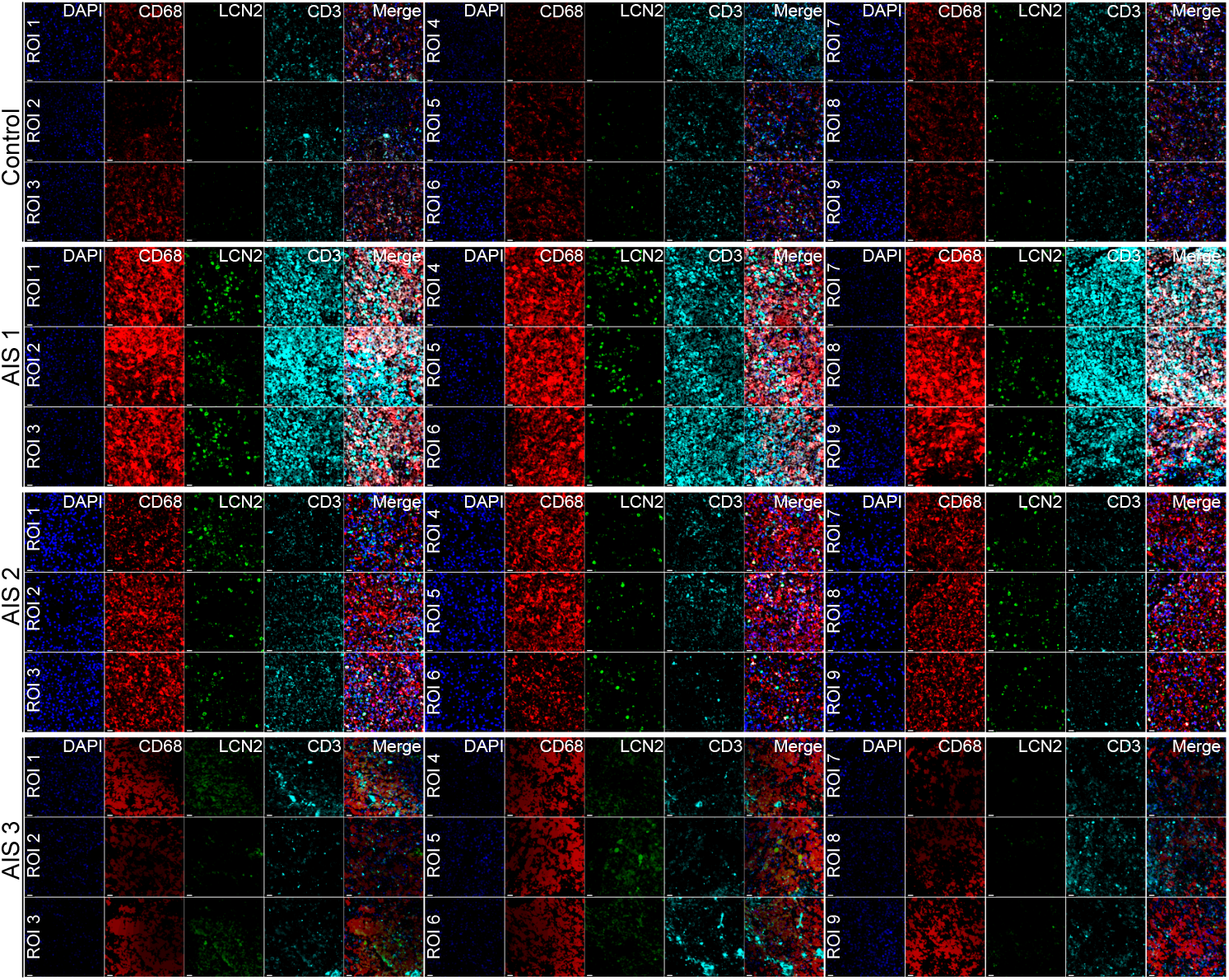
– related to Figure 5. Lipocalin-2 expression by CD68^+^ red pulp macrophages is conserved in human ischemic stroke spleens. The figure displays the various regions of interest (ROIs) that were used for quantitative image assessment shown in Figure 5G in one reference non-stroke control spleen as opposed to three biologically distinct spleens analyzed from patients with acute ischemic stroke (AIS). Scale bar: 20 µm.

**Supplementary Figure 7.**
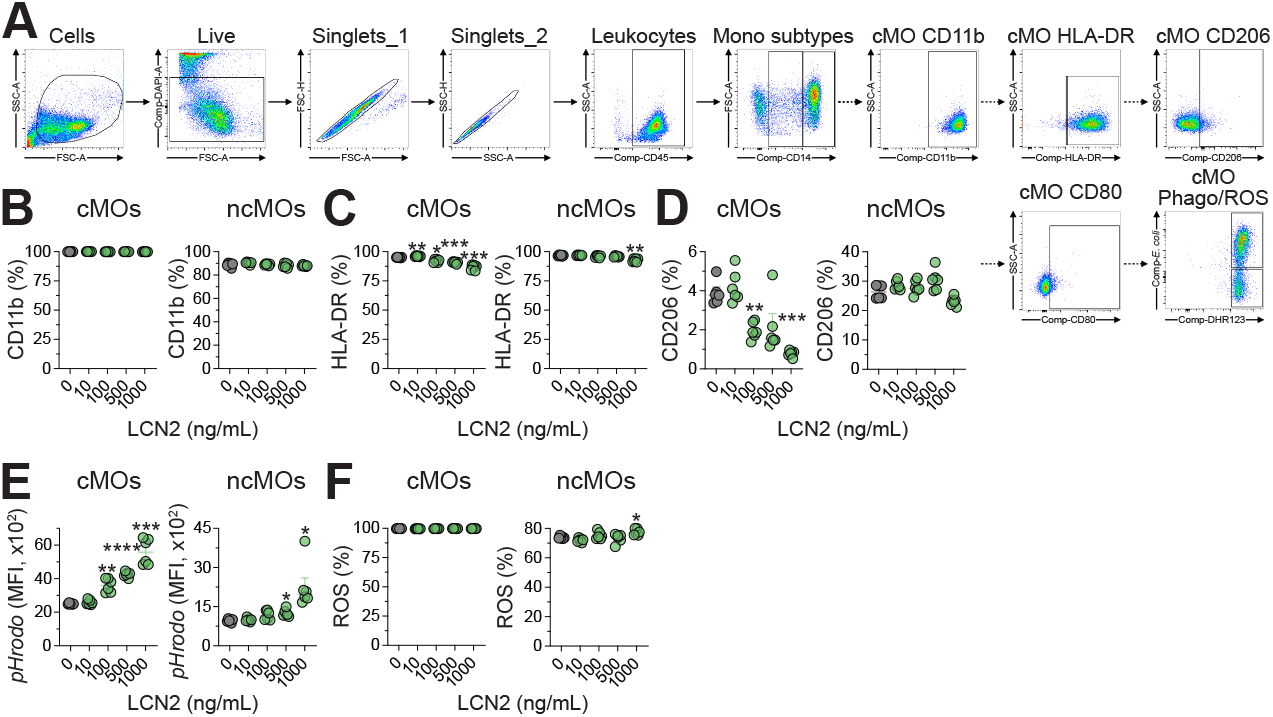
– related to Figure 6. Exposure to recombinant lipocalin-2 is linked to immunomodulatory priming in human monocytes. (**A**) Gating strategy for flow cytometry to identify cells by FSC-A and SSC-A to exclude debris followed by identification of live cells (DAPI negative) and sequential selection of singlets (singlets_1: FSC-A vs. FSC-H; singlets_2: SSC-A vs. SSC-H). Next, leukocytes were identified by CD45 positivity followed by monocyte subtype identification based on CD14 expression. Classical monocytes (cMOs) were CD14^HI^ vs. non-classical, i.e., intermediate monocytes were CD14^DIM^ (ncMOs). Both cMOs and ncMOs were used to characterize their expression levels of CD11b, HLA-DR, CD80, CD206 or their ability to phagocytose *E*. *coli* bioparticles or produce reactive oxygen species (ROS). (**B**) Percentage of CD11b on cMOs (left) or ncMOs (right), respectively, within untouched monocytes after stimulation with vehicle– or LCN2 for 4 hours assessed by flow cytometry. (**C-D**) Same as (**B**) but HLA-DR (**C**) or CD206 (**D**). (**E**) Phagocytic competence as indicated by mean fluorescence intensity (MFI) of pHrodo^+^ cMOs (left) or ncMOs following a 30-minute exposure to pHrodo-labeled *E*. *coli* bioparticles (25 µg/mL). (**F**) Production of ROS (in %) during exposure to *E*. *coli* bioparticles in cMOs (left) or ncMOs (right). (**B-F**) Data (*n* = 6 technical replicates per group) are median ± IQR and representative of two independent experiments (biological donors). Statistical comparisons were performed with a one-way ANOVA, with Tukey’s multiple comparison test for parametric data. \**P* ≤ 0.05, \*\**P* ≤ 0.01, \*\*\**P* ≤ 0.001, \*\*\*\**P* ≤ 0.0001.

## Materials and methods

### Availability of data and materials

#### Lead contact

Further information and requests for resources or materials reported in this study should be directed to and will be fulfilled by the corresponding author.

#### Data and code availability

The dataset supporting the conclusions of this article is included within the article (and its additional file(s)). Differential expression results obtained from 3’-MACE-Seq are provided as supplementary tables (Supplementary Table 1 and 2). Image-analysis scripts and processed quantification of files are available from the corresponding author upon reasonable request.

### Ethics declarations

#### Ethics approval and consent to participate

All experimental procedures were performed in strict accordance with the protocols approved by the Goethe University Frankfurt am Main or as set out by the local committees. The use of human samples was approved by the Ethical Review Committee of the Goethe University Frankfurt am Main (approval no. 329/10 (use of human PBMCs from German Red Cross), 19-232 (“Investigation of the role of sphingosine-1-phosphate receptor 1 modulation as a mediator of stroke-induced lymphopenia”; DRKS 00033903), and 4/09 (project number SGI-13-2018; use of human non-stroke spleen)). The use of human spleens from stroke patients (n = 3) was approved by the local Ethics Review Committee at Charité (approval number EA1/144/13) and the Ethical Review Committee of the Goethe University Frankfurt am Main (approval number 119/26). Autopsy consent was obtained from the patients or their families. Identities were encoded at the Department of Neuropathology before sharing for sample processing or data analysis. All human research conformed to the Declaration of Helsinki and the Department of Health and Human Services Belmont Report.

#### Consent for publication

Written informed consent was obtained from all participants/legal guardian(s) included in the study. The authors confirm: (1) that the work described has not been published before (except in the form of an abstract or as part of a published lecture, review, or thesis); (2) that it is not under consideration for publication elsewhere; (3) that its publication has been approved by all co-authors, if any; (4) that its publication has been approved (tacitly or explicitly) by the responsible authorities at the institution where the work is carried out.

### Mice

All animal procedures in the present study conformed to the German Protection of Animals Act and guidelines for care and use of laboratory animals by the local committee (Regierungspräsidium Darmstadt, Germany, FU/1143). For all experiments, male C57BL/6J mice, 10-12-weeks-old, obtained from Charles River Laboratories (Sulzfeld, Germany) were used and housed on a 12/12-hour light-dark cycle with food and water *ad libitum*.

### Transient middle cerebral artery occlusion and post-operative care

We performed transient focal cerebral ischemia by intraluminal occlusion of the right middle cerebral artery (tMCAO) as described previously (<u>Lucaciu et al. 2020</u>) (84). Briefly, mice were anesthetized with 1.5% isoflurane in oxygen. Following a midline cervical incision, the right common carotid artery (CCA), external carotid artery (ECA), and internal carotid artery (ICA) were exposed. After ligation of the CCA and ECA, a silicone rubber-coated monofilament (6-0 medium; tip diameter 0.23 mm; 6023910PK10, Doccol Corporation, Sharon, MA, USA) was introduced into the ICA and advanced to occlude the origin of the MCA. After 60 min of occlusion, the filament was carefully withdrawn to allow reperfusion. In sham-operated controls, the filament was inserted to the level of the MCA origin and immediately withdrawn without sustained occlusion. We continuously monitored body temperature, which was maintained at 37°C ± 0.5°C using a heating mat throughout surgery and recovery. Pre-emptive analgesia was administered by intraperitoneal injection of buprenorphine (0.1 mg/kgBW) 30 min before surgery and continued every 8 hours for 48 hours post-intervention. Postoperative care for mice was provided as described previously (<u>Kestner et al. 2020</u>) (120). Animals were housed in groups of 3-5 mice per cage on a heating pad, and post-stroke care involved regular monitoring protocols, including daily assessment of body weight, rectal temperature, and scoring of neurological deficits. For mice subjected to 3 days or 7 days stroke endpoint, softened pellet food, jelly food, and standard pellets were placed on the cage floor from day 2 onwards to facilitate easy access to food. Animals received oral food supplementation (2-3 mL dissolved chow per day) and subcutaneous injections of 0.9% warm saline (2-4 mL/day). Neurological deficits were assessed after 24 hours, 3 days, and 7 days post-surgery using the functional Experimental Stroke Scale (ESS) (<u>Lourbopoulos et al. 2017</u>) (121). Induction of AIS as a result of tMCAO was confirmed by 2,3,5-triphenyltetrazolium chloride (TTC) staining as described previously (<u>Popp et al. 2009</u>) (122).

### Exclusion Criteria and Mortality

The following criteria were used for exclusion of animals from subsequent analysis: 1. Death during surgery, 2. Subarachnoid hemorrhage (SAH) detected upon euthanasia, 3. Absence of AIS following TTC staining.

In total, 51 mice were used in this study. Of these, all were randomly assigned to the study groups. Here, 10 were allocated to the 24 hour post-tMCAO timepoint, 17 to the 3 days post-tMCAO timepoint, 16 mice to the 7 days post-tMCAO timepoint, and 8 mice were allocated to sham treatment. Treatment specific mortality was 10% (1 of 10; 24 hours post-tMCAO), 23.5% (4 of 17; 3 days post-tMCAO), and 25% (4 of 16; 7 days post-tMCAO), respectively. No sham-operated animal died. Accordingly, 9, 13, 12, and 8 animals, respectively were available for analysis as only surviving animals were included.

Spleens were processed as described below and divided into three portions (RNA, protein, and FFPE), allowing the same animals to contribute to multiple downstream readouts. Consequently, the group sizes reported in individual figure legends refer to animals contributing to the respective assay and are not additive across assays.

Samples undergoing immunofluorescence, RT-qPCR, or Western blot analysis were provided in a blinded manner. Unblinding was performed after analysis was completed.

### 3’-MACE-Sequencing

#### Library preparation and sequencing library preparation (MACE_D, MACE-Seq)

The 3’-RNA libraries were prepared using the MACE-Seq Kit (GenXPro GmbH) according to the manufacturer’s instructions. Fragmentation of mRNA was performed, the 3’ ends were reverse transcribed, and cDNA was generated after template switching and incorporation of Unique Molecular Identifiers (UMIs, TrueQuant; GenXPro), followed by PCR amplification with a minimal number of cycles and silica-bead-based purification.

#### Sequencing MACE_D, MACE-Seq

Sequencing was performed on an Illumina NEXTSEQ instrument with 1×76 bps.

#### Raw data processing

Raw data processing is a critical first step in a sequencing analysis, ensuring high-quality input samples used for downstream applications. Two essential components of this process are trimming and UMI deduplication. Trimming removes low-quality bases, sequencing adapters, and other technical artifacts from raw reads to enhance alignment accuracy, using tools such as Trimmomatic or Cutadapt. UMI deduplication is particularly important in experiments using UMIs to tag individual molecules prior to amplification. By collapsing reads with identical UMIs and inserts, in-house developed tools were used to effectively eliminate PCR duplicates, reducing amplification bias and improving quantification accuracy. Together, trimming and UMI deduplication were used to refine the dataset to ensure more reliable results for gene expression profiling and variant detection. Unprocessed sequencing reads were adapter-trimmed and quality-trimmed using Cutadapt 4.6. UMI deduplication was done using in-house developed tools collapsing PCR duplicates using UMIs into single reads. Using Cutadapt 4.6 additional sequencing artifacts were removed from the reads in an extra cleaning step. FastQC 0.11.9 was used to assess the quality of sequencing reads.

### Spleen collection

At defined experimental time points, mice were euthanized by cervical dislocation in accordance with institutional guidelines. The spleen was rapidly excised and divided into three portions. One piece was immediately snap-frozen in liquid nitrogen for RNA isolation, one piece was used for protein extraction, and the third piece was fixed in 4% formalin for histological analysis.

### RNA isolation, cDNA synthesis, and real-time, quantitative polymerase chain reaction

#### RNA isolation

One of the excised pieces of the spleen was immediately transferred to Eppendorf tubes together with Tri reagent (Sigma-Aldrich, T9424, Taufkirchen, Germany), homogenized and total RNA was isolated according to the manufacturer’s instructions. RNA purity and concentration were determined using NanoDrop (ThermoFisher, Darmstadt, Germany). The A_260/280_ ratio for all samples was ≥ 1.9.

#### cDNA synthesis

A total of 1 µg of RNA was reverse transcribed into cDNA using the RevertAid Reverse Transcriptase kit (Thermo Scientific, K1621, Darmstadt, Germany) according to the manufacturer’s instructions.

#### Real-time, quantitative polymerase chain reaction

RT-qPCR was performed using the TaqMan Gene Expression Assay in 96-well PCR plates (Applied Biosystems, Thermo Fisher Scientific, Darmstadt, Germany) and run in a Quantstudio3 PCR system (Applied Biosystems, Thermo Fisher Scientific, Foster City, CA, USA). Relative gene expression levels were calculated using the ΔΔCT method by normalizing fold-change of expression to an endogenous housekeeping gene (*GAPDH*). The results are presented as the fold-change of expression relative to control samples. All Taqman primers purchased from LifeTechnologies (ThermoFisher Scientific, Darmstadt, Germany) used in this study are as follows: *Gapdh* (Mm99999915_g1), *Lcn2* (Mm01324470_m1), *Il6* (Mm00446190_m1), *Il1b* (Mm00434228_m1), *Tnf* (Mm00443258_m1).

### Western blotting

The snap-frozen spleen samples were homogenized in protein lysis buffer (M-PER, 78503, Thermo Fisher) supplemented with protease and phosphatase inhibitors and centrifuged at 13,000 rpm for 15 minutes at 4°C. Supernatants were collected and total protein amounts were determined using the BCA Assay Kit (23227, Thermo Fisher, Schwerte, Germany) according to the manufacturer’s instructions. 25 µg of protein lysates were resolved by SDS polyacrylamide gel electrophoresis (SDS-PAGE) and transferred onto a 0.2 µm PVDF membrane (03010040001, Roche, Mannheim, Germany) at 70 mA per gel for 75 minutes at room temperature. The membranes were blocked in 2.5% non-fat milk in TBS-T and incubated in primary antibodies overnight at 4°C. The primary antibodies used in this study include anti-β-actin (1:25,000 dilution, A2228, Sigma Aldrich) and anti-Lipocalin-2 (1:1,000 dilution, AF1857, R&D Systems). Following overnight incubation, the primary antibody was washed off, and membranes were incubated with HRP-conjugated secondary antibodies (1:10,000 dilution), anti-mouse IgG (SA00001-8) and anti-goat IgG (SA00001-3) (all Proteintech, Rosemont, IL, USA) for 60 minutes at room temperature. Protein bands were visualized using enhanced chemiluminescence (Pierce^TM^ ECL, 32106, Thermo Fisher Scientific) and imaged with an iBright^TM^ 1500 system (Thermo Fisher Scientific).

### Immunofluorescence and bioinformatic analysis

#### Immunofluorescence

Formalin fixed, paraffin-embedded spleen tissues were prepared in 4 µm-thick sections, deparaffinized in xylene, rehydrated through a series of alcohols to distilled water. Targeted epitopes were unmasked by heat-induced epitope retrieval (HIER) at 100°C for 20 minutes in citrate buffer (S1699, Dako). The sections were permeabilized with 0.1% Triton X-100 for 5 minutes and blocked with 1× Rotiblock blocking solution (A151.1, Roth) for 30 minutes at room temperature. The sections were incubated at 4°C overnight with primary antibodies at a dilution of 1:200. The antibodies used in this study include anti-F4/80 (28463-1-AP, Proteintech), anti-Lipocalin-2 (AF1857, R&D Systems), anti-CD3 (CL488-17617, Proteintech), anti-CD3 (sc1127, Santa Cruz Biotechnology), anti-CD68 (CL-488-28058, Proteintech), anti-CD68 (754068882, Invitrogen), and anti-Ly6G (BE0075-1, BioXCell). Following overnight incubation, fluorescently-labeled antibodies were applied for 1 hour at room temperature in the dark. Nuclei were counter-stained with DAPI (D9542, Sigma) at 1 µg/mL for 60 seconds and sections were mounted in Dako fluorescent mounting medium (S3023, Dako). Images were acquired using a Keyence BZ-X fluorescence microscope under identical exposure settings for comparative analyses.

The use of human material was approved by the local Ethics Committee of the Faculty of Medicine of the University Hospital Frankfurt (Voting-No 4/09, approval year 2009, project number SGI-13-2018). Quantification of marker expression by immunofluorescence of human non-stroke control or spleens obtained from patients with ischemic stroke was performed using QuPath with standardized thresholding and segmentation parameters applied uniformly across all images.

#### Image processing and quantitative analyses

Image processing and quantitative analyses were performed in Python 3.11 using custom scripts based on NumPy, scikit-image, and OpenCV (<u>Harris et al. 2020</u>; <u>Van Der Walt et al. 2014</u>; <u>Bradski 2000</u>) (123–125). Image intensities were manually adjusted before image export. Exported fluorescence images were converted to grayscale intensity images, smoothed using a Gaussian filter (5 x 5 kernel), and converted to 8-bit intensity values. Local contrast was enhanced using contrast-limited adaptive histogram equalization (CLAHE; clip limit 2.0, tile grid size 20 x 20).

For murine spleen analyses, marker-positive regions were segmented using multi-level Otsu thresholding (four classes), with the highest intensity threshold used for binary segmentation of positive staining. For CD3 staining, a minimum threshold intensity of 70 was applied to limit background classification as positive staining in images containing scarce CD3 signals. Red-pulp regions of the spleen were manually delineated in overview fluorescence images using napari (v0.6.4) (<u>Sofroniew et al. 2026</u>) (126). The annotations were exported as binary masks and applied to the segmented CD3, F4/80, and LCN2 channels to restrict quantification to red pulp regions. Two to three images were analyzed per mouse.

For human spleen analysis, 9 red pulp regions of interest (ROIs) were manually selected per patient. These ROIs exhibited stronger fluorescence signals and different intensity distributions from the murine overview images; therefore, a separate thresholding approach was selected. Marker-positive regions were segmented using an image-specific Li threshold, rounded to the nearest integer and constrained to a minimum intensity of 5.

Marker-positive area was quantified as the number of positive pixels divided by the reference area in pixels, multiplied by 100. The reference area was the mask-defined red pulp area for murine images and the entire red-pulp ROI for human images. CD3-, F4/80-, and LCN2-positive areas were quantified in murine samples or CD68– and LCN2-positive areas in human samples. In addition, pixel-level double-positive area was determined by intersecting the binary LCN2 mask with the F4/80 mask in murine samples or the CD68 mask in human samples, respectively. The resulting overlap was expressed as a percentage of the reference area.

Following image export, preprocessing parameters and threshold-selection rules were applied consistently across experimental groups within each species-specific workflow without sample-specific manual adjustment. For quality control, validation images were generated for each analyzed image or ROI, showing the original fluorescence channels alongside the masked binary segmentations for murine samples and with colored segmentation overlays for human samples.

### Mouse Inflammation Antibody Array

Inflammatory proteins in spleen lysates were analyzed using the Mouse Inflammation Antibody Array (ab133999, Abcam) according to the manufacturer’s instructions. In brief, snap-frozen spleen tissues were homogenized in lysis buffer supplemented with protease inhibitors. After centrifugation at 13,000 *g* for 15 minutes at 4°C, supernatants were collected. Total protein concentrations were determined by the BCA Assay (23227, Thermo Fisher, Schwerte, Germany). 100 µg of protein lysate for each sample were incubated with the provided array membranes, which were pre-spotted with capture antibodies against 40 inflammatory cytokines. After over-night incubation at 4°C on a gentle shaker, membranes were washed and incubated with a biotinylated detection antibody cocktail, followed by HRP-conjugated streptavidin. Subsequently, signals were visualized using enhanced chemiluminescence (ECL) and detected with an iBright^TM^ Imaging System (Thermo Fisher Scientific). Densitometric analysis was performed using the ImageJ software. Signal intensities were background-corrected and normalized to the positive control spots on each membrane to account for inter-membrane variability. Relative protein expression levels were calculated and expressed as fold change compared to control samples.

### Splenocyte and magnetic-bead-based immune cell isolation

Male, C57BL/6J mice, aged 10-12 weeks, were euthanized and spleens isolated as described before. Spleens were aseptically harvested and transferred into ice-cold RPMI1640 medium (11875093, Gibco) without serum. Single cell suspensions were prepared by gentle mechanical dissociation, followed by filtration through a 70 µm cell strainer (352350, Corning) into 15 mL conical tubes. Cells were washed once with RPMI1640 medium and centrifuged at 500 *g* for 5 minutes at 4°C. The supernatant was discarded and red blood cells were lysed using 1x RBC Lysis Buffer (420301, Biolegend) for 10 minutes at 4°C. The reaction was stopped by adding excess RPMI1640 medium, and cells were centrifuged at 500 *g* for 10 minutes at 4°C. The resulting splenocytes were resuspended in FACS buffer (PBS containing 0.5% BSA and 2 mM EDTA) for magnetic separation. Using the Pan T Cell Isolation Kit, T cells and myeloid cells were subsequently enriched using magnetic-bead-based separation according to the manufacturer’s protocol (130-096-535, Miltenyi). Briefly, 1×10^7^ splenocytes were incubated with anti-biotin microbeads for 10 minutes at 4°C. After washing, labeled cells were separated using LS columns (130-042-401, Miltenyi). The positive fraction (pan T cells) and negative fraction (myeloid cells) were collected separately and used for further downstream applications.

### Transwell Migration Assay

Chemotactic migration of isolated T cells was investigated using a 96-well transwell system consisting of 5.0 µm-sized pores in a polycarbonate membrane (3388, Corning). A total of 7.5×10^4^ T cells were seeded in the upper chamber in 100 µL RPMI1640 medium containing 0.1% BSA. The lower chamber contained 200 µL migration medium (RPMI1640 supplemented with 10 mM HEPES and 0.1% BSA) with or without 200 ng/mL recombinant CCL5 protein (250-07, Peprotech) and in combination with 0-500 ng/mL recombinant LCN2 protein (50060-MNAH, SinoBiological). After incubation for 5 hours at 37°C and 5% CO_2_, cellular migration to the lower chamber was assessed by counting the migrated cells in the lower chamber using a hemocytometer.

### Flow cytometric assessment of phagocytosis-induced ROS production

Single-cell suspensions of murine splenocytes were collected as described before and resuspended in RPMI1640 medium supplemented with heat-inactivated fetal bovine serum (FBS), 1% penicillin/streptomycin, and 1% L-glutamine. Cells were counted and seeded at 1×10^5^ splenocytes per well in 200 µL medium in 96-well Nunclon Sphera Ultra Low Attachment Plates (Thermo Fisher Scientific). All cultures were maintained at 37°C in a humidified incubator with 5% CO_2_. Splenocytes were then stimulated for 4 hours with vehicle or recombinant murine LCN2 protein (1857-LC-050, R&D Systems/Bio-Techne, Minneapolis, USA) at final concentrations of 0, 10, 100, 500, or 1000 ng/mL. Following stimulation, unbound LCN2 protein was washed off and cells were immediately processed for functional assays. Phagocytic activity was assessed using DEEP red pHrodo-labeled *E*. *coli* bioparticles (Thermo Fisher Scientific) according to the manufacturer’s instructions. Prior to addition to the culture medium, bioparticles were opsonized by incubation with 10% heat-inactivated AB serum for 30 minutes at 37°C. Opsonized bioparticles were added to each well at a final concentration of 25 µg/mL, and plates were incubated for 30 minutes at 37°C, 5% CO_2_. To assess phagocytosis-induced ROS production, 1 µM of dihydrorhodamine 123 (DHR-123, Thermo Fisher Scientific) was supplemented to the cocktail of *E*. *coli* bioparticles. After 30 minutes of phagocytosis and ROS production, the cells were washed with FACS buffer and kept on ice until preparation for flow cytometry. Here, after three washes and centrifugation in FACS buffer (400 *g*, 5 minutes, 4°C), splenocytes were incubated with murine FcX TrueStain Fc block (BioLegend) for 10 minutes at 4°C to minimize non-specific binding. Surface staining was performed for 60 minutes in the dark at 4°C on ice in the presence of murine FcX TrueStain Fc block using the following fluorochrome-conjugated antibodies (all BioLegend): Ly6C (BV785, clone HK1.4), F4/80 (BV650, clone BM8), CD45 (BV421, clone 30-F11), CD11b (PE/Cy7, clone M1/70), Ly6G (PE, clone 1A8), CD19 (APC/Cy7, clone 6D5), and CD3 (APC/Cy7, clone 17A2). DAPI (BioLegend) was included for live/dead discrimination. After staining, splenocytes were washed and resuspended in FACS buffer. Samples were then acquired on a BD LSR Fortessa II flow cytometer (BD Biosciences) using standard instrument settings and compensation controls. Data were analyzed with FlowJo software (BD Biosciences). Gating strategies included the exclusion of debris and doublets, selection of live CD45^+^ leukocytes, and identification of myeloid and lymphoid subsets based on established marker combinations. All functional assays were performed in ≥ three technical replicates and ≥ two independent experiments.

### Human PBMC and magnetic-bead-based monocyte isolation

Peripheral blood samples from healthy adult donors were obtained as buffy coats from the German Red Cross, with written informed consent and approval from the local Ethics Committee (#329/10, Goethe University Frankfurt). Peripheral blood mononuclear cells (PBMCs) were isolated by density gradient centrifugation using Ficoll-Paque Plus according to standard protocols, ensuring high yield and viability of mononuclear cells. Following isolation, untouched monocytes were enriched from PBMCs using the Pan Monocyte Isolation Kit (Miltenyi Biotec) and LS columns, following the manufacturer’s instructions. This negative selection approach routinely yields monocyte purities of ≥ 95% with high viability, while minimizing activation and preserving functional integrity. Purity and viability were routinely assessed by cytospin preparations and trypan blue staining. Additional washes were performed to minimize platelet contamination, in line with current best practices for monocyte isolation (<u>Nielsen et al. 2020</u>) (101).

### Flow cytometric assessment of phagocytosis-induced ROS production in untouched human monocytes

For functional assays, monocytes were seeded at 1×10^5^ cells per well in 96-well Nunclon Sphera Ultra Low Attachment plates (Thermo Fisher Scientific) in RPMI-1640 medium supplemented with 10% heat-inactivated human AB serum, 1% penicillin/streptomycin, and 1% L-glutamine. Cells were pre-stimulated for 4 hours with recombinant human LCN2 protein (1757-LC-050, R&D) at concentrations of 0, 10, 100, 500, or 1000 ng/mL, reflecting physiologically relevant and pathological ranges observed under states of acute inflammation (<u>Ni et al. 2013</u>) (127). Control wells received vehicle only. Phagocytic activity was assessed using DEEP red pHrodo-labeled *E*. *coli* bioparticles (Thermo Fisher Scientific), which were opsonized with 10% heat-inactivated human AB serum for 30 minutes at 37°C prior to addition to the culture medium. Opsonization with human serum enhances physiological uptake by monocytes via Fc and complement receptors, maximizing assay sensitivity and specificity (<u>Kapellos et al. 2016</u>) (128). Opsonized bioparticles were added to each well at a final concentration of 25 µg/mL, and plates were incubated for 30 minutes at 37°C, 5% CO_2_. To assess phagocytosis-induced ROS production, 1 µM of DHR-123 (Thermo Fisher Scientific) was supplemented to the cocktail of *E*. *coli* bioparticles. After 30 minutes of phagocytosis and ROS production, the cells were washed with FACS buffer and kept on ice until preparation for flow cytometry. Here, after three washes and centrifugation in FACS buffer (400 *g*, 5 minutes, 4°C), monocytes were first incubated with human Fc block (BioLegend) for 10 minutes at 4°C to minimize non-specific antibody binding. Surface staining was performed in the presence of human Fc block for 60 minutes at 4°C in the dark on ice using fluorochrome-conjugated antibodies: DAPI (live/dead discrimination), HLA-DR (BUV395, clone L243, BD Biosciences), CD206 (BV605, clone 15-2, BioLegend), CD11b (BV421, clone M1/70, BioLegend), CD14 (PE/Cy5, clone TüK4, Invitrogen), CD80 (PE-CF594, clone 2D10, BioLegend), and CD45 (PE, clone HI30, BioLegend). After staining, cells were washed and resuspended in FACS buffer for acquisition. Samples were acquired on a BD LSR Fortessa II flow cytometer using standardized instrument settings and compensation controls. Gating strategies included exclusion of debris and doublets, selection of live CD45^+^ leukocytes, and identification of classical (CD14^high^) vs. intermediate monocytes (CD14^dim^). All functional assays were performed in ≥ three technical replicates and ≥ two independent experiments.

### Statistical analyses

Data distribution was assessed for normality using the Shapiro-Wilk test. Depending on the outcome, data are presented as mean ± SD for normally distributed variables or a median ± IQR for non-normally distributed variables. *N* is defined separately in each figure legend and refers to either biologically independent samples (individual mice or human donors) or to technical replicates, as specified. For *ex vivo* stimulation experiments, *n* denotes technical replicates derived from a defined number of biologically independent animals or donors, which is stated in the respective legend.

#### Two-group comparisons

Non-normally distributed data were compared using the Mann-Whitney U test. For paired experimental designs in which cells derived from the same animal or human donor were allocated to vehicle and LCN2 pre-conditioning, a paired two-tailed t test was applied.

#### Multi-group comparisons

For normally distributed data, a one-way ANOVA was used. Where all groups were compared against a single reference condition (vehicle or sham), Dunnett’s multiple comparisons test was applied; where all pairwise comparisons were of interest, Tukey’s multiple comparisons test was applied. For repeated-measures designs in which all treatment conditions were derived from the same biological sample, a repeated-measures one-way ANOVA with Geisser-Greenhouse correction was used, followed by Dunnett’s multiple comparisons test. For non-normally distributed data, the Kruskal-Wallis test followed by Dunn’s multiple comparisons test was used; for matched non-parametric designs, the Friedman test followed by Dunn’s multiple comparisons test was applied.

#### Reporting of significance

A two-sided *P* value < 0.05 was considered statistically significant. Exact *P* values are reported in the figure legends where a comparison approached but did not reach this threshold. Statistical analyses were performed using GraphPad Prism (v9) and R (v4.5.0). Transcriptomic analyses were carried out in R; image-derived quantifications were generated in Python 3.11 and subsequently analyzed in GraphPad Prism.

## Supporting information

Full Table 2

## Acknowledgments

The authors are most grateful to the individuals and their relatives for consenting to autopsy and subsequent research, which were facilitated by the Biobank of the Department of Neuropathology, Charité – Universitätsmedizin Berlin and the German National Autopsy Network (NATON – 01KX2524). This work is supported by the German Research Foundation (SFB1039-TPB08), the Leducq Foundation (SphingoNet), and the Uniscientia Stiftung, Vaduz. A. Lucaciu was funded by the Patenschaftsmodell of the Frankfurt-Forschungs-Förderung. J.S. was funded by the Deutsche Forschungsgemeinschaft (Clinician Scientist position, SU 1360/1-1). S.S. and J.S. are participants of the national Translational Tandem Program for Gene– and Cell-based Therapies (nTTP-GCT) – coordinated by the Biomedical Innovation Academy of the Berlin Institute of Health at Charité (BIH) and funded by the Federal Ministry of Education and Research (BMBF). Open Access funding was enabled and organized by Projekt DEAL. Illustrations were created with BioRender.com under a commercial license (graphical abstract, Fig. 1A, Fig. 3A/C, Fig. 4A, Fig. 5A, Fig. 6A). Language editing and grammar refinement were supported by an AI-based tool (you.com). The authors reviewed and edited all AI-assisted text and take full responsibility for the scientific content.

## Authors’ contributions

Lucaciu, J. Subburayalu, and R. Vutukuri conceived the project and designed the experiments. A. Lucaciu, P. Wurzel, S. R. Rasmussen, E. Lueckhoff, F. Mayser, R.-I. Kestner, V. Haas, L. S. Huber, D. Bevara, J. Subburayalu, and R. Vutukuri performed the experiments. P. Wurzel and J. Subburayalu performed the bioinformatic analyses. A. Lucaciu, P. Wurzel, J. Subburayalu, and R. Vutukuri interpreted the data. J. Benjamin and S. Subramanian provided technical support. N. Landvogt, M. Glueck, K. Gertz, R. Raspe, C. Welsch, J. Bein, P. J. Wild, H. Radbruch, C. Grefkes, A. Strzelczyk, W. Pfeilschifter, M. H. Sieweke, and J. M. Pfeilschifter provided critical reagents and resources. P. Wurzel and R. Vutukuri contributed to writing the manuscript. A. Lucaciu and J. Subburayalu wrote the manuscript. All authors read, discussed, and commented on the manuscript.

---

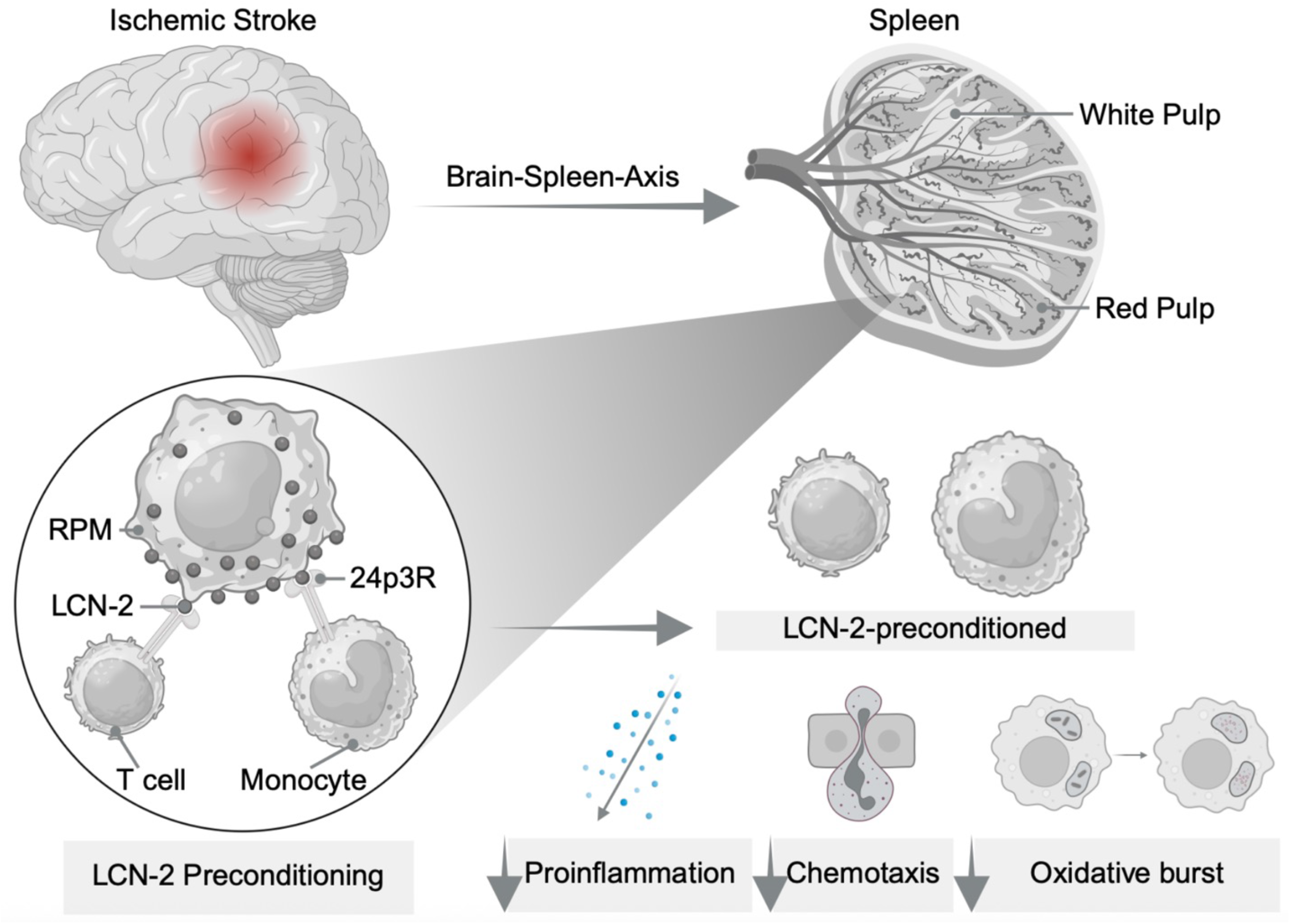

